# The binary protein interactome mapping of the *Giardia lamblia* proteasome lid reveals extra proteasomal functions of GlRpn11

**DOI:** 10.1101/2024.09.18.613619

**Authors:** Ankita Das, Atrayee Ray, Nibedita Ray Chaudhuri, Soumyajit Mukherjee, Shubhra Ghosh Dastidar, Alok Ghosh, Sandipan Ganguly, Kuladip Jana, Srimonti Sarkar

**Affiliations:** Department of Biological Sciences, Bose Institute, Unified Academic Campus, EN 80, Sector V, Bidhannagar, Kolkata - 700091, West Bengal, India; Department of Biochemistry, University of Calcutta, 35 Ballygunge Circular Road, Kolkata - 700019, West Bengal, India; Division of Parasitology, National Institute for Cholera and Enteric Diseases, P33- C.I.T. Road, Scheme XM, Beleghata, Kolkata - 700010, West Bengal, India

**Keywords:** *Giardia*, Proteasome, Mitosome, GlRpn11, GlRpn8

## Abstract

*Giardia lamblia* does not encode Rpn12 and Sem1, two proteins crucial for assembling the proteasome lid. To understand how the interactions between the giardial proteasome lid subunits may have changed to compensate for their absence, we used the yeast two-hybrid assay to generate a binary protein interaction map of the *Giardia* lid subunits. Most interactions within the *Giardia* proteasome lid are stronger than those within the *Saccharomyces cerevisiae* lid. These may compensate for the absence of Rpn12 and Sem1. A notable exception was the weaker interaction between GlRpn11 and GlRpn8, compared to the strong interaction between Rpn11-Rpn8 of yeast. The Rpn11-Rpn8 dimer provides a platform for lid assembly and their interaction involves the insertion of a methionine residue of Rpn11 into a hydrophobic pocket of Rpn8. Molecular modeling indicates that GlRpn8’s pocket is wider, reconciling the experimental observation of its weak interaction with GlRpn11. This weaker interaction may have evolved to support extra proteasomal functions of GlRpn11, which localizes to multiple subcellular regions where other proteasome subunits have not been detected. One such location is the mitosome. Functional complementation in yeast shows that GlRpn11 can influence mitochondrial function and distribution. This, together with its mitosomal localization, indicates that GlRpn11 functions at the mitosome. Thus, this parasite’s proteasome lid has a simpler subunit architecture and structural attributes that may support dual functionalities for GlRpn11. Such parasite-specific proteasome features could provide new avenues for controlling the transmission of *Giardia*.

^1^

**Graphical Abstract:** 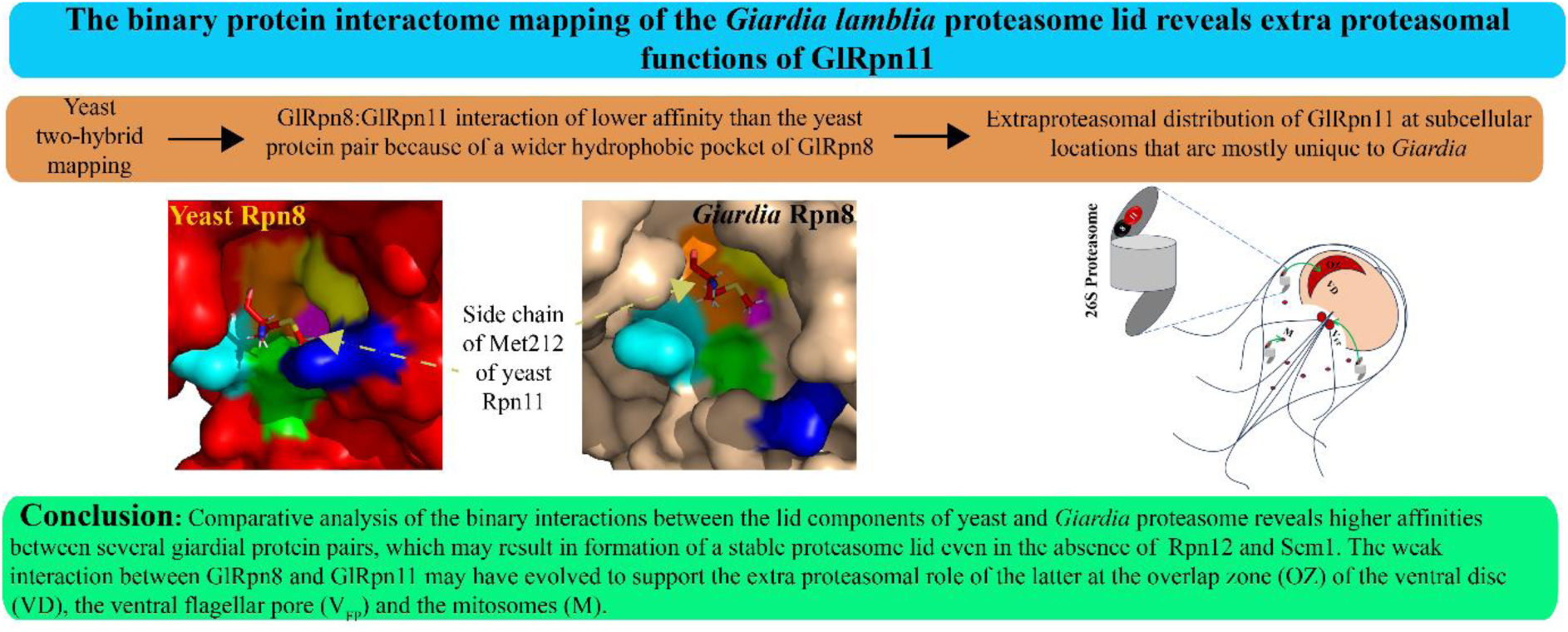

*Highlights:*

- *Giardia* genome does not encode two proteasomal lid subunits: Rpn12 and Sem1
- Unique interactions within the lid may compensate for the absence of these two
- GlRpn8:GlRpn11 weakly interacts to support GlRpn11’s extra-proteasomal distribution
- GlRpn11 localizes at mitosomes, OZ of VD, and to the V_FP_
- The 182-218 fragment of GlRpn11 may regulate mitosomal function

## 1. Introduction

*Giardia lamblia* is a unicellular flagellated pathogen that parasitizes the host gut to cause the diarrheal disease giardiasis. The parasite has two morphologically different lifecycle stages: the pear-shaped trophozoites and the oval cysts. While trophozoites are the disease-causing form, the metabolically quiescent cysts are infective as they can survive the inhospitable conditions present outside the host. Cysts ingested by the host undergo excystation within the host gut. Trophozoites that emerge colonize the upper part of the small intestine and multiply by binary fission [1]. If these trophozoites are washed further down the alimentary tract, cyst formation occurs via encystation. Cysts are released into the environment by the host excretion process [1]. Thus, both encystation and excystation are vital for the survival and transmission of this parasite.

Given the difference in morphological and biochemical properties of these two forms, it is expected that their proteomes would also differ [2]. Changes in the proteome composition involve both protein synthesis and degradation. Studies in model organisms have established that cellular protein degradation is carried out by either the proteasome or the lysosome [3]. The proteasome is a large, 2.5 MDa multi-subunit complex that harbours multiple proteases having broad substrate specificity [4]. It is an integral part of all eukaryotic cells as it turns over proteins that are either dysfunctional or no longer required. It originated before the advent of eukaryotes as large protease complexes have been observed in archaea and some eubacteria [5]. The proteasome degrades proteins in both the cytosol and the nucleus in a ubiquitin-dependent manner [6,7]. Each proteasome comprises a 20S core particle (CP) and a 19S regulatory particle (RP) that caps either one or both ends of the CP. The CP harbours protease activity. The RP has two subcomplexes- a base proximal to the CP and a lid distal to the CP. Studies of the RP from model organisms indicate that the base contains six AAA+ ATPase subunits (Rpt1-6) and three non-ATPase subunits (Rpn1, Rpn2 and Rpn13), whereas the lid is composed of nine non-ATPase subunits (Rpn3, Rpn5, Rpn6, Rpn7, Rpn8, Rpn9, Rpn11, Rpn12, and Sem1) [8]. Another non-ATPase subunit, Rpn10, resides at the lid-base interface [8]. The functions of the RP include recognition of ubiquitylated substrates by two proteasomal ubiquitin receptors, Rpn10 and Rpn13 [9,10], followed by deubiquitination by Rpn11, and finally, substrate unfolding by the base ATPases [13].

Curiously, giardial orthologues of Rpn12, Rpn13, and Sem1 could not be identified by either sequence-based searches of the genome or mass spectrometric analysis [12,13]. Since both Rpn13 and Rpn10 are ubiquitin receptors, the latter may compensate for the former’s absence [14]. However, the absence of Rpn12 and Sem1 is intriguing as these two proteins play seminal roles in the assembly of the lid, which proceeds via the formation of intermediates. One intermediate comprises Rpn5, Rpn6, Rpn8, Rpn9 and Rpn11. These together constitute Module 1, in which a stable heterodimer of Rpn8-Rpn11 is joined by the other three [8]. Rpn3 and Rpn7 form a separate intermediate, lid particle 3 (LP3), tethered by Sem1, also known as Rpn15 [15]. Module 1 associates with LP3, creating LP2. The last subunit, Rpn12, completes the assembly process as it triggers a large-scale conformational remodeling of LP2 that ultimately drives lid assembly [16,17]. Therefore, the absence of Rpn12 and Sem1 in the *Giardia* genome indicates that the quaternary structure of its proteasomal lid is likely to be different from that of its opisthokont hosts. This hypothesis is further supported by sequence analyses. *Giardia* encodes all the base subunits of the RP, most of which share at least 40% sequence identity with the corresponding yeast orthologue (Supplementary Table 1). Hence, it is expected that their assembly will be similar to that of yeast. On the contrary, the putative giardial lid subunits are more diverged from their yeast orthologues, with the identity values of most subunits being around 20%. Since proteasomal inhibition is known to cause the production of nonviable *Giardia* cysts, this prospect of an altered lid architecture may lead to the designing of new anti-*Giardia* molecules that specifically inhibit the lid assembly process in this human pathogen [18].

To understand the molecular organization of the lid subunits of the *Giardia* proteasome, we have generated a protein-protein interaction map of these subunits using yeast two-hybrid (Y2H) analysis. Such an approach has been previously used to determine the proteasome assembly in *S. cerevisiae*, *C. elegans*, and *H. sapiens* [19–21]. Many of these interactions are in agreement with the subsequent structural studies [22,23]. Our results indicate a considerable difference in the binary interaction profile between the lid subunits of *Giardia* and those of yeast. Comparative analysis of interaction affinities between orthologous protein pairs from yeast and *Giardia* indicates that while many giardial subunits display stronger affinities, some binary protein interactions are weaker in this protist, notably in the GlRpn8-GlRpn11 pair. A possible underlying reason for this weaker interaction could be that GlRpn11 discharges proteasome-independent functions at multiple locations within *Giardia* trophozoites. Its localization revealed that apart from the expected nuclear and cytoplasmic distribution, GlRpn11 is also present at mitosomes, the overlap zone (OZ) of the ventral disc (VD) and the ventral flagellar pores (V_FP_) of trophozoites. Extraproteasomal role is also documented for yeast Rpn11 as it regulates mitochondrial function [24]. Our results show that GlRpn11 can functionally substitute for the yeast Rpn11 as it can not only function as a proteasomal deubiquitinase but can also modulate yeast mitochondrial function. This ability of GlRpn11, together with its localization to mitosomes, raises the possibility that it may be involved in regulating the mitosomal function in *Giardia*.

## 2. Materials and methods

### 2.1. Bioinformatic analysis

The protein sequences of *Giardia* proteasomal subunits were retrieved from GiardiaDB (www.giardiadb.org) by BLAST search analysis using sequences of proteasome lid component orthologues from *Saccharomyces cerevisiae* and *Homo sapiens* as query. Functional domain analyses were performed using SMART. Secondary structure predictions were carried out using Phyre^2^ [25] and AlphaFold [26]. Based on such predictions, multiple sequence alignment was performed in ClustalW [27] followed by editing and visualization using Jalview [28]. The conserved residues and those at the contact surfaces were mapped in PyMOL using the crystal structure of Rpn8-Rpn11 heterodimer (PDB ID: 4O8X) [29] of *S. cerevisiae* as a template.

### 2.2. Culture of <u>Giardia lamblia</u> trophozoites and induction of encystation

*G. lamblia* (strain ATCC 50803/ WB clone C6) trophozoites were grown in TYI-S-33 media (pH 6.8) supplemented with 10% adult bovine serum and 0.5 mg/ml bovine bile [30]. Upon reaching confluency, encystation was induced as previously described [31].

### 2.3. Yeast two-hybrid assay

For creating bait (fusion with the DNA binding domain of Gal4) and prey (fusion with the activation domain of Gal4) constructs, the *Giardia* and yeast orthologues of *RPN3*, *RPN5*, *RPN6*, *RPN7*, *RPN8*, *RPN9* and *RPN11* were PCR amplified using primers listed in Supplementary Table 2. The PCR products were cloned in the bait vector, pGBT9 (*TRP1* selection marker), and the prey vector, pGAD424 (*LEU2* selection marker) (Clonetech Laboratories), using specific restriction enzymes whose sites were incorporated in the primers (Supplementary Table 2). Different pairwise combination of these bait and prey constructs was co-transformed into PJ69-4A strain and selected on yeast complete medium (YCM) plates lacking leucine and tryptophan (LT). The interaction between the BD and AD fusion proteins was monitored by assessing the expression of the *ADE2*, *HIS3,* and *LacZ* reporter genes. *HIS3* expression was assessed by monitoring growth on YCM plates lacking leucine, tryptophan, and histidine but containing 2.5 mM 3-anino-1,2,4-triazole (LTH 3-AT), and *ADE2* expression was assessed on YCM plates lacking leucine, tryptophan, and adenine (LTA). Plates were incubated at 28°C for three days. The β-galactosidase activity was estimated by colorimetry, and the values are the average of readings obtained from three independent experiments, with two technical replicates for each sample. The results were statistically validated using a two-tailed unpaired t-test in GraphPad Prism 5, where a *p*-value of <0.01 was considered statistically significant.

### 2.4. Raising polyclonal antibodies against GlRpn11 and GlTom40 followed by determination of antibody specificity by western blot analysis

*glrpn11* and *gltom40* were PCR amplified from *G. lamblia* genomic DNA with primers listed in Supplementary Table 2 and cloned in pET32a (Novagen). The His-tagged fusion proteins were expressed in *E. coli* BL21 (DE3). GlTom40 expression was induced for 4 h at 37°C, and GlRpn11 expression was induced for 16 h at 20°C. GlTom40 was isolated from the pellet fraction, and GlRpn11 was isolated from the supernatant. The purified proteins were used to raise polyclonal antibodies in rat (GlTom40) or rabbit (GlRpn11). All animal experiments adhered to the guidelines are approved by the Institutional Animal Ethical Committee of Bose Institute (IAEC/BI/136/2019). The specificity of each antibody was determined by performing western blotting with trophozoite protein extract. The images of uncropped membranes are presented in Supplementary Data 1.

### 2.5. Immunofluorescence

Immunofluorescence was performed in trophozoites and encysting trophozoites as described previously [32]. Briefly, cells were harvested by chilling the tubes on ice for 20 min, followed by centrifugation at 1000 g for 10 min and washed twice with 1X PBS. Cells were fixed for 15 min with 4% paraformaldehyde in 1X PBS at RT. After fixation, the cells were collected by centrifugation and treated with 0.1 M glycine for 8 min at RT. Permeabilization was performed in trophozoites and encysting trophozoites with 0.1% Triton X-100 (v/v) in 1X PBS for 6 min at RT, followed by blocking with 2% BSA for 2 h at RT. Cells were incubated at 4°C overnight with preimmune sera, anti-GlRpn11, or anti-GlTom40, in 1:50 dilution in 0.2% BSA in 1X PBS solution. The cells were harvested by centrifugation and washed thrice with 1X PBS (10 min each wash), followed by incubation for 2 h in the dark with 1:400 dilutions of either goat-anti-rabbit Alexa-Fluor 594 conjugated secondary antibody or goat-anti-rat Alexa-Fluor 488 conjugated secondary antibody. The cells were washed thrice with 1X PBS, re-suspended in antifade reagent (0.1% of p-phenylene-diamine in 90% glycerol solution) and mounted on the slide. Samples were observed with a confocal laser scanning microscope (Leica Stellaris 5) using a 63X objective. Images were assembled with Adobe Photoshop CS3 and Adobe Illustrator CS3.

### 2.6. Site-directed mutagenesis

Mutations were introduced using KOD Hot Start DNA Polymerase (EMD Merck Millipore) as per the manufacturer’s recommended PCR conditions by using overlapping primers [33] to PCR-amplify the entire construct bearing the respective gene of interest. The PCR product was digested with *Dpn*1 (NEB) at 37°C for 2 h and transformed into *E. coli* DH5α strain. Plasmids were isolated, and the presence of the desired mutation was confirmed by sequencing. Primers and constructs are listed in Supplementary Table 2 and Supplementary Table 3.

### 2.7. Functional complementation assay in S. cerevisiae

The assay was carried out in the BY4742 strain. The essential genes *RPN8* and *RPN11* were deleted using plasmid shuffling [34]. Briefly, the yeast genes were cloned into *URA3* selectable marker containing vector, using primers listed in Supplementary Table 2. These constructs were transformed into BY4742. Next, the chromosomal copy of either *RPN8* or *RPN11* was deleted in transformants carrying episomal copies of the gene to be deleted. The chromosomal deletions were performed with PCR products generated using gene-specific primers (Supplementary Table 2), with the *HIS5Mx6* deletion cassette as a template [35]. The deletions were confirmed via PCR using specific primers listed in Supplementary Table 2. For plasmid shuffling, *glrpn8* and *glrpn11* were individually cloned into a *LUE2* selectable marker containing vector and transformed into the corresponding yeast deletion mutant, *rpn8Δ*/*rpn11Δ*. Next, these double transformants were grown on YCM liquid medium lacking leucine so that the cells growing in the presence of uracil may be cured of the *URA3* construct bearing the yeast gene. Plasmid loss was assessed by selection on YCM plates containing 0.5% 5-fluoroorotic acid to ensure the loss of the episomal copies of the yeast genes. The chromosomal gene deletions were once again confirmed via PCR, as mentioned above. The strains and constructs generated are listed in Supplementary Table 3.

### 2.8. Measurement of oxygen consumption rate

To measure whole cell respiration rate, yeast cells were grown in liquid medium and harvested at the late log phase (OD_600_ = 1), followed by resuspension in fresh 2 % glycerol-containing media at a concentration of 10^8^ cells/ml. The oxygen consumption rate (OCR) was measured at 30°C using an Oyxtherm fitted with a Clack Electrode (Hansatech, UK). Respiration was measured for 4 min and blocked by adding 1 mM sodium azide. The average OCR (nmol of oxygen produced/min/10^8^ cells) was estimated from the data of at least three biological replicates (n = 9). The baseline and the nonspecific OCR (after sodium azide addition) were negligible. The results were statistically validated using a two-tailed unpaired t-test in GraphPad Prism 5, where a *p*-value of <0.01 was considered statistically significant.

### 2.9. Molecular dynamics simulation

PDB files of both Rpn8 and GlRpn8 were retrieved from AlphaFold [26]. The initial coordinates were processed in the CHARMM-GUI web server [36]. The models were solvated in a water box, ensuring at least 10 Å TIP3P [37] water layer everywhere, while maintaining 0.15 M KCl concentration. The systems were energy minimized briefly, followed by a short gradual phase of heating up to 300 K and a 1 ns equilibration of the systems. All-atom molecular dynamics simulations were next performed at a constant temperature of 300 K and 1 atm pressure [38] (i.e., NPT ensemble) using the CHARMM36 force field [39] and NAMD 2.13 package [40]. A 2 fs time step was used for numerical integration of the dynamics while the hydrogen bonds were constrained using the SHAKE algorithm [41]. A short-range non-bonded cut-off of 12 Å was used, and the PME method [42] was used to compute the long-range electrostatics, under periodic boundary conditions. A simulation of 20 ns was done to relax and optimize each system. The minimum distances between three of the rim residues of Rpn8 from yeast and *Giardia* were measured in each frame across the simulated period of 20 ns, and the data corresponding to the last 15 ns were collected and processed for further analysis. The change in distance (Å) of the three rim residue pairs from both the proteins over the last 15 ns were plotted.

## 3. Results

### 3.1. Interaction between the Rpn9 and Rpn5 orthologues is conserved between yeast and Giardia

One of the intermediate subcomplexes in the proteasome lid assembly process is Module 1, comprising the PCI domain-containing subunits Rpn9, Rpn5 and Rpn6, and the MPN domain-containing subunits Rpn8 and Rpn11 [8]. The interactions between the PCI domains result in an Rpn9-Rpn5-Rpn6 trimer that interacts with the Rpn8-Rpn11 dimer. The PCI-PCI interaction surface involves a hydrophobic core surrounded by electrostatic interactions along the periphery [22]. Since orthologues of all Module 1 subunits are encoded in the *Giardia* genome, we wanted to understand if the interactions between them are conserved in this protist. Based on reports of strong interaction between Rpn5 and Rpn9 by Y2H, as well as biochemical experiments, we studied the interaction between GlRpn5 and GlRpn9 to see if a similar interaction is preserved [19,43]. Multiple sequence alignment shows that the hydrophobic and charged residues that contribute to the PCI-PCI interaction surface in the yeast Rpn9-Rpn5 pair are largely conserved in the *Giardia* orthologues (Supplementary Fig. 1).

The *Giardia* and yeast orthologues were expressed as fusions of the Gal4 DNA binding domain (BD) or the Gal4 activation domain (AD). We expressed different pairwise combinations of the BD and AD-fused proteins in the yeast host strain and observed the expression of *HIS3* and *ADE2* in transformants with AD-Rpn5 and BD-Rpn9. Expression of the highly stringent *ADE2*, in addition to the less stringent *HIS3* [44], indicates a high affinity interaction between the two yeast proteins (Fig. 1A). Transformants with AD-GlRpn5 and BD-GlRpn9 fusions did not turn on the expression of even *HIS3*. However, quantitative estimation of the *LacZ* reporter activity indicated a possible weak interaction between the two *Giardia* proteins (Fig. 1B). Since incompatibility with AD and/or BD is known to cause false-negative results in Y2H [45], we repeated the assay after interchanging the domains. AD-GlRpn9 and BD-GlRpn5 interacted strongly as all three reporter genes were activated (Fig. 1A, B). Notably, the yeast pair AD-Rpn9 and BD-Rpn5 did not interact in this orientation. To confirm that even though there is a difference in the orientation between the yeast and *Giardia* pair, the interaction between AD-GlRpn9 and BD-GlRpn5 is still mediated by the PCI domains, we mutated Glu343 in AD-GlRpn9. This residue aligns with Glu325 of Rpn9, a known key contributor to the electrostatic interactions that form part of the PCI-PCI interaction between Rpn9 and Rpn5 [43]. When BD-GlRpn5 was expressed with AD-GlRpn9* (Glu343Lys), none of the three reporter genes were expressed (Fig. 1A, B). This indicates that the interaction between GlRpn9 and GlRpn5 is similar to that between the orthologous yeast protein pair. Thus, even though GlRpn5 and GlRpn9 share low sequence identity with their yeast counterparts (18.9% for Rpn5 and 18.5% for Rpn9), the nature of the interaction between these two protein partners is similar to that in yeast.

**Fig. 1.**
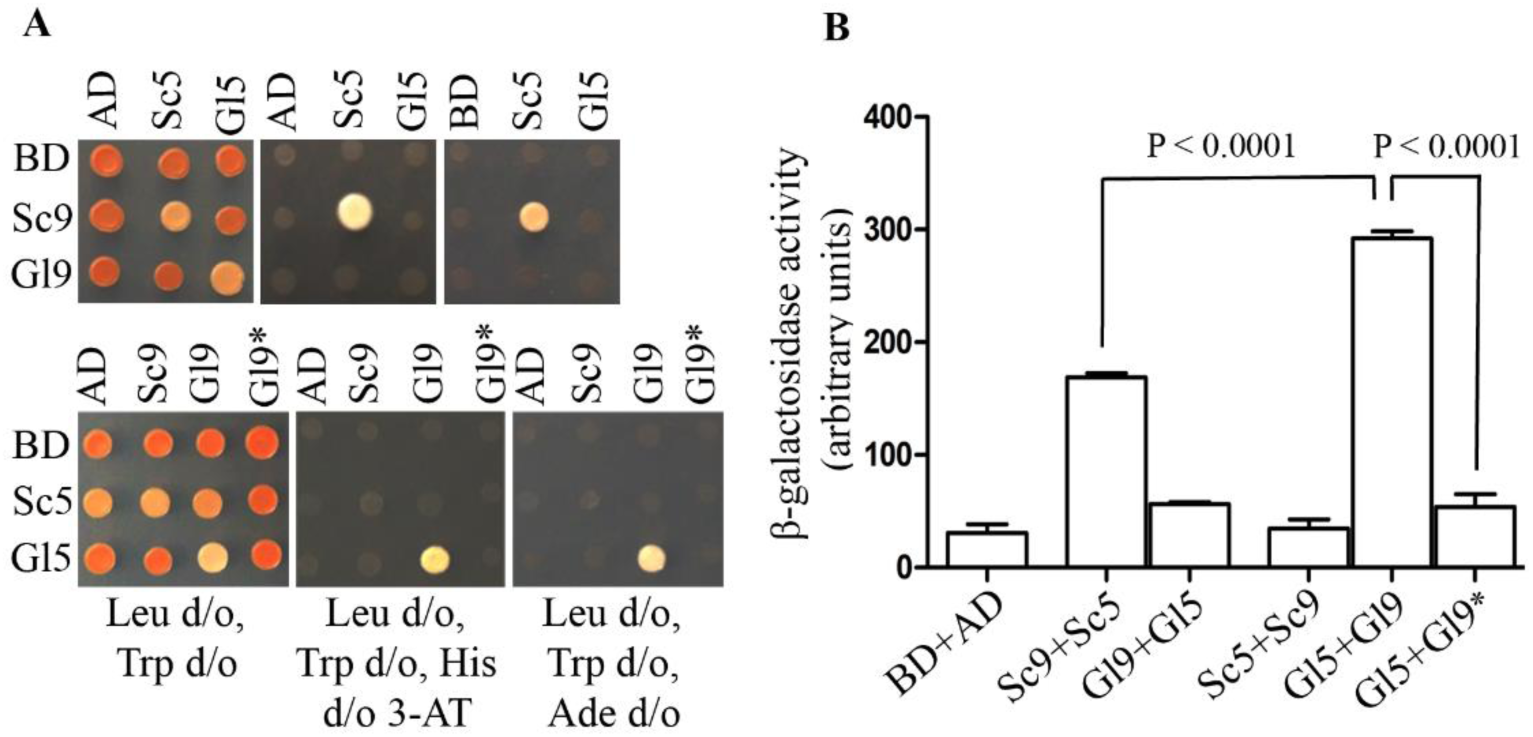
Interaction between the Rpn9 and Rpn5 orthologues is conserved between yeast and *Giardia*. A) PJ69-4A strain was transformed with combinations of constructs expressing *Saccharomyces cerevisiae* or *Giardia lamblia* orthologues of Rpn5 (Sc5/Gl5) or Rpn9 (Sc9/Gl9) as fusions of the Gal4 activation domain (AD) or Gal4 DNA binding domain (BD), as indicated. The interaction between the various fusion combinations was assessed by *HIS3* and *ADE2* reporter gene expression by monitoring the growth of the transformants on plates lacking leucine, tryptophan, and histidine with 2.5 mM 3-AT (Leu d/o, Trp d/o, His d/o 3- AT), or lacking leucine, tryptophan, and adenine (Leu d/o, Trp d/o, Ade d/o), respectively. Growth on plates lacking leucine and tryptophan (Leu d/o, Trp d/o) indicates the presence of the fusion constructs. All plates were incubated at 28°C for three days. B) Quantification of β-galactosidase activity of transformants expressing various combinations of AD and BD fusions as described in (A) above. The statistical significance of the difference in the interaction between two interacting protein pairs is represented as follows: *** *p*-value *<* 0.0001; ** *p*-value *<* 0.001; * *p*-value *<* 0.01; ns *p*-value *>* 0.01 where n = 6. The raw data underlying this graph is given in Supplementary Data 2.

### 3.2. Multiple strong binary interactions within the proteasome lid of Giardia

The Rpn3-Rpn7 interaction plays a crucial role in the proteasomal lid assembly process in yeast. Together with Sem1, they form the intermediate LP3 [15]. To compensate for the absence of Sem1 in *Giardia*, we hypothesized that the interaction between GlRpn3 and GlRpn7 is likely to be different than that in yeast. While Fu et al observed that the BD-Rpn3 could autoactivate the reporter genes, they documented interaction between Rpn3 and Rpn7 using AD-Rpn3 [19]. Consistently, we, too, observed autoactivation by BD-Rpn3 as it alone turned on both *HIS3* and *ADE2* (Fig. 2A). Unlike BD-Rpn3, there was no autoactivation with BD-GlRpn3, which indicates differences in the surface distribution of amino acids between the two orthologues. The known Rpn3-Rpn7 interaction in yeast was observed with AD-Rpn3 and BD-Rpn7, which turned on both reporter genes (Fig. 2A). In contrast, the interaction between BD-GlRpn7 and AD-GlRpn3 was weak, as only *HIS3* was turned on (Fig. 2A). These results indicate that the interaction between GlRpn3 and GlRpn7 has changed compared to that between the orthologous pair from yeast.

**Fig. 2.**
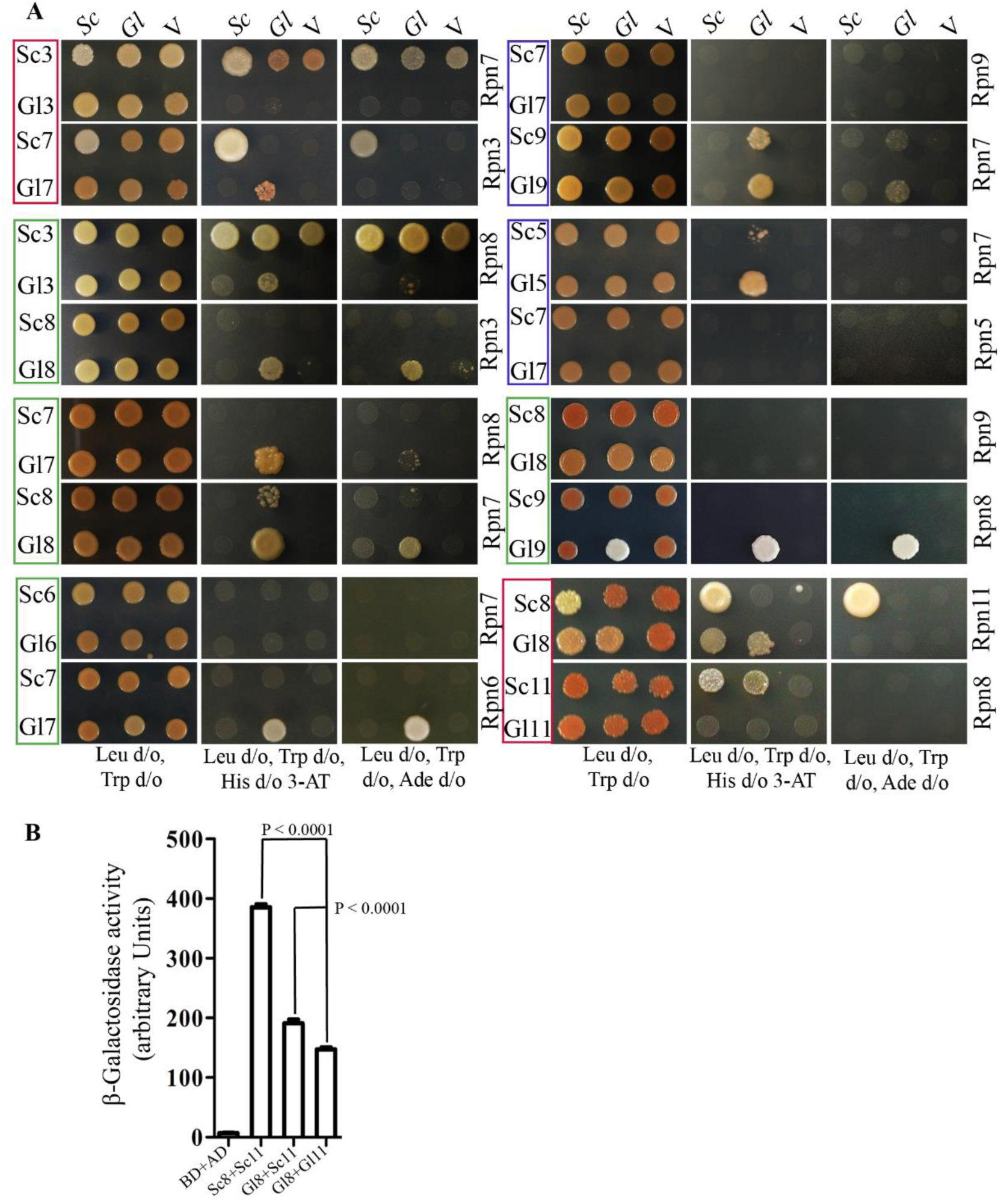
Multiple strong protein-protein interactions within the proteasome lid of *Giardia*. A) The interactions between various proteasome lid subunits of *S. cerevisiae* (Sc) and *G. lamblia* (Gl) were assessed as described in Fig.1A. Proteins with the BD domain fused to their N-terminus are indicated on the left of each panel (Sc3, Gl3, etc.). The name on the right of each panel (Rpn3, Rpn7, etc.) indicates the lid subunit whose yeast (*Sc*) or *Giardia* (*Gl*) orthologue (as indicated above the panels) are expressed as fusions having the AD domain at their N-terminus. *V* dictates empty vector pGAD424. Green boxes denote those protein pairs where the *Giardia* orthologues exhibit higher affinity, compared to the corresponding ones in yeast. Red boxes denote interactions stronger in the yeast orthologues than those between the corresponding *Giardia* proteins; blue boxes mark pairs with weak interactions between the *Giardia* protein pairs, compared to the lack thereof between the yeast orthologues. Combinations where no interaction was detected for either the yeast or *Giardia* pairs are shown in Supplementary Fig. 2. B) Quantification of β-galactosidase activity to compare the interaction between Rpn8-Rpn11 and GlRpn8-GlRpn11. β-galactosidase activity was estimated in transformants expressing various combinations of AD and BD fusions, as described in (A). The statistical significance of the difference in the interaction between two interacting protein pairs is represented as follows: *** *p*-value *<* 0.0001; ** *p*-value *<* 0.001; * *p*-value *<* 0.01; ns *p*-value *>* 0.01, where n = 6. The raw data underlying this graph can be found in Supplementary Data 3.

The weak interaction between GlRpn3 and GlRpn7 may be compensated by each of these two subunits having stronger interactions with other lid subunits. Although the previous exhaustive Y2H study did not detect any interaction between yeast Rpn3 and Rpn8 [19], another high throughput functional study in yeast documented synthetic interactions between them [46]; the cryo-EM structure of the yeast proteasome also shows physical contact [22,47]. Therefore, we tested if GlRpn3 has any affinity for GlRpn8. Consistent with Fu et al. [19], we failed to detect any interaction between Rpn3 and Rpn8 of yeast (Fig. 2A). But GlRpn3 interacts strongly with GlRpn8 as evidenced by the growth in the absence of not only histidine but also adenine. GlRpn8 also interacted strongly with GlRpn7, but the corresponding interaction was not detected between Rpn8 and Rpn7 (Fig. 2A), which, too, share physical contact in the cryo-EM structure of the yeast lid [47]. The same applies to the strongly interacting GlRpn7-GlRpn6 pair (Fig. 2A), whose yeast orthologues are nearby in the cryo-EM structure, but they fail to interact [47]. GlRpn7 also interacts, albeit weakly, with both GlRpn5 and GlRpn9 (Fig. 2A). Thus, even though GlRpn3 and GlRpn7 interact weakly, each of them interacts strongly with other lid subunits, and this may enable their incorporation into the *Giardia* proteasome lid even in the absence of Sem1. Our data indicates that in contrast to Rpn8, GlRpn8 interacts strongly not only with GlRpn3 and GlRpn7 but also with GlRpn9 (Fig. 2A). Thus, unlike in yeast, the lid of *Giardia* appears to be stabilized through multiple strong interactions of GlRpn8, *viz.* with GlRpn3, GlRpn7, and GlRpn9. Additional unique interactions of GlRpn7 with GlRpn6, GlRpn5, and GlRpn9 may contribute to this stabilization.

### 3.3. GlRpn8 and GlRpn11 are functionally orthologous to the corresponding yeast proteins but have lower affinity for each other than those of yeast

Considering the strong interactions of GlRpn8 with multiple lid subunits, we assessed its interaction with GlRpn11. As previously mentioned, the Rpn8-Rpn11 dimer is a part of Module 1 [8]. Consistent with the previous Y2H study, we observed a strong interaction between BD-Rpn8 and AD-Rpn11 as cells grew in the absence of histidine and also adenine (Fig. 2A) [19]. In contrast, BD-GlRpn8 and AD-GlRpn11 interacted weakly with growth in the absence of histidine only. This was also supported by quantitative estimation of β-galactosidase activity as cells expressing the yeast protein pair had nearly double the activity compared to those expressing the *Giardia* proteins (Fig. 2B). The weak interaction between GlRpn8 and GlRpn11 is unusual given that of all the lid subunits, these two share the highest sequence identity with the corresponding yeast orthologues, 28% for GlRpn8 and 40.4% for GlRpn11 (Supplementary Table 1). Such weak interaction raises the possibility that these proteins may not be proteasomal subunits, rather their paralogues as the proteasome lid subunits are known to bear close homology to components of other cellular complexes, the COP9 signalosome and the eukaryotic translation initiation faction, eIF3 [48]. In yeast, the JAB1/MPN/MOV34 metalloenzyme domain is present not only in Rpn8 and Rpn11 but also in Rri1 and Prp8. While the former is an isopeptidase of the catalytic subunit of the COP9 signalosome and cleaves Nedd8, the latter is a component of U4/U6-U5 snRNP complex [49,50]. To determine if the putative GlRpn8 and GlRpn11 are indeed functionally orthologous to Rpn8 and Rpn11, we carried out functional complementation in yeast with the premise that since both *RPN8* and *RPN11* genes are essential, survival of the yeast deletion mutant strains expressing the corresponding *Giardia* gene will indicate that the yeast and *Giardia* proteins are functional orthologues. The *Giardia* genes were introduced in the respective yeast deletion mutants (*rpn8Δ* or *rpn11Δ*) strains by the plasmid shuffling process as described previously [51]. Briefly, chromosomal deletions were carried out using the *HIS5* cassette in the presence of the respective yeast genes on a *URA3* vector. The corresponding *Giardia* gene expressed from a *LEU2* vector was introduced. Next, the yeast genes were shuffled out by selecting against the *URA3* marker using 5-fluoro-orotic acid.

Both the *rpn8Δ* and *rpn11Δ* strains bearing the *glrpn8* and *glrpn11* genes, respectively, lost the yeast orthologues as indicated by the failure of these cells to grow on media lacking uracil (Fig. 3A, B). Growth on media lacking leucine indicates the presence of the giardial gene, while growth media lacking histidine indicates that the chromosomal copies of the two essential genes, *RPN8* or *RPN11*, are absent. The survival of these cells indicates that the giardial orthologue of the deleted gene was able to functionally substitute for the corresponding yeast gene (*RPN8* and *RPN11*). The presence of the yeast, instead of the giardial orthologues on the *LEU2* vector, served as positive control.

**Fig. 3.**
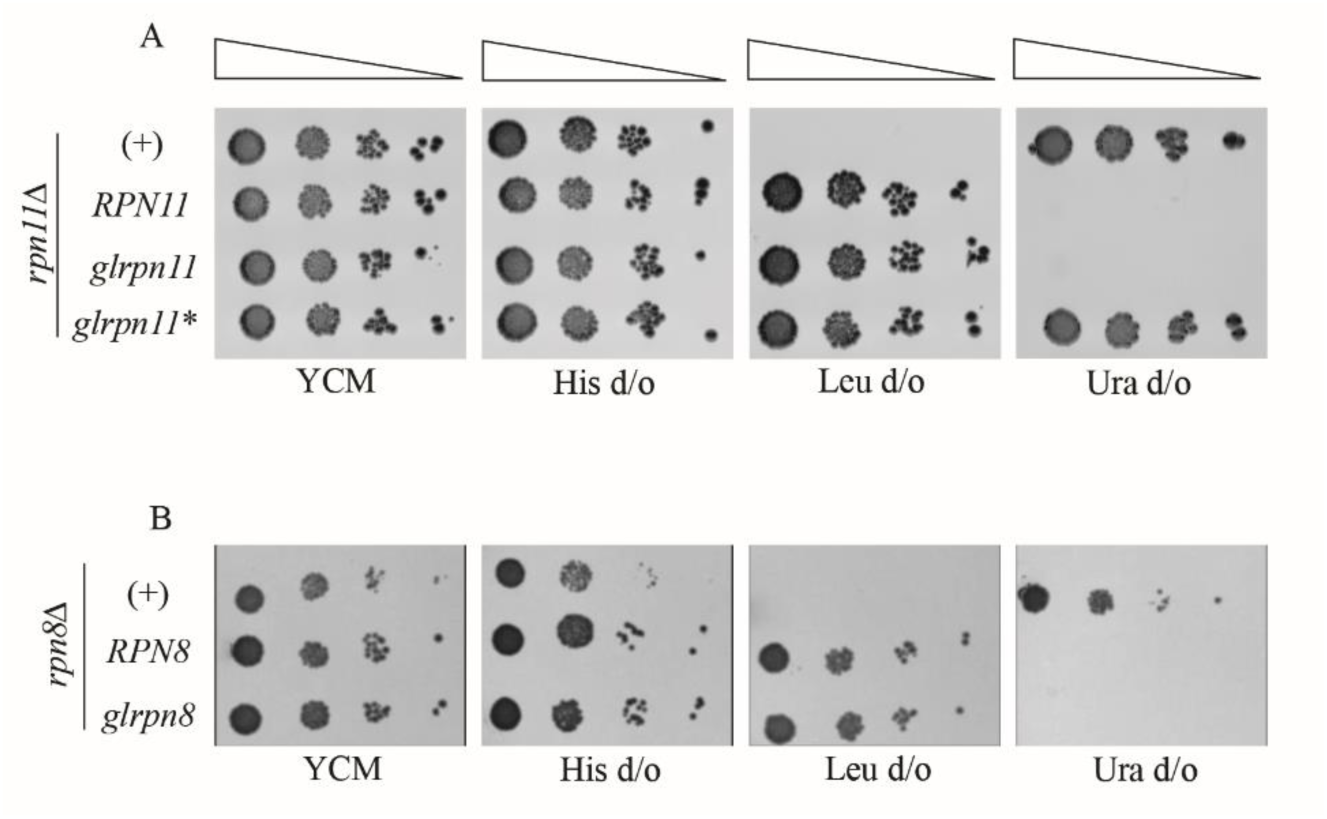
GlRpn11 and GlRpn8 can functionally substitute for the corresponding orthologous proteins from yeast. A) Expression of GlRpn11 allows survival of yeast cells with chromosomal deletion of the essential gene *RPN11*. The growth of *rpn11Δ* mutant expressing Rpn11, GlRpn11 or mutant GlRpn11* (His120Ala, His122Ala), from episomal vector, on complete medium (YCM), synthetic defined medium lacking histidine (His d/o), or leucine (Leu d/o), or uracil (Ura d/o). Rpn11 was expressed from a *URA3* marked vector (+) or *LEU2* marked vector (*RPN11*). GlRpn11 and GlRpn11* are expressed from a *LEU2* marked vector. B) Expression of GlRpn8 allows the survival of yeast cells with chromosomal deletion of the essential gene *RPN8*. Same as (A), except *RPN8* or *glrpn8* is expressed episomally, as indicated.

Both Rpn11 and Rpn8 contain an MPN domain at their N-terminal end. But this domain in Rpn11 has the MPN+ motif that makes it a functional deubiquitinase [52]. The absence of this motif in Rpn8 renders it catalytically inactive [52]. The signature sequence of the MPN+ is HXHX_7_SX_2_D. These two His residues bind to a Zn^2+^ ion, which is necessary for the catalytic activity of Rpn11. His➔Ala mutations are lethal in yeast [11]. Our analysis shows the presence of an MPN+ motif in the sequence of GlRpn11. To explore if the two His residues present within the MPN+ of GlRpn11 (His120 and His122) are also functionally significant, site-directed mutagenesis was used to replace them with Ala. The *rpn11Δ* strain, expressing this double mutant (GlRpn11*), was unable to functionally substitute for the yeast protein (Fig. 3A). Thus, similar to the Rpn11, an MPN+ motif is also essential for the deubiquitination function of GlRpn11. Incidentally, the sizes of the colonies expressing Glrpn8 or Glrpn11 in *rpn8Δ* or *rpn11Δ*, respectively, were similar to those expressing the corresponding yeast orthologues from the *LEU2* plasmid. This proves that not only are the parasite proteins orthologous to the yeast proteins, but their presence in the yeast proteasome does not cause any detectable perturbations in proteasomal function. Both Rpn11 and GlRpn11 could not functionally substitute for Rpn8, and neither could the Rpn8 and GlRpn8 do the same for Rpn11, indicating that similar to yeast, GlRpn8, and GlRpn11 perform unique functions in the context of the proteasome, even if both of them contain the MPN domain (data not shown). Taken together, the results of these experiments indicate that GlRpn8 and GlRpn11 are functionally orthologous to the corresponding yeast proteins (Fig. 3A, B).

We investigated why GlRpn8 and GlRpn11 interact weakly. The crystal structure of the Rpn8-Rpn11 heterodimer (PDB ID: 4O8X) shows that these proteins interact via their MPN domains [29]. This dimerization involves two interfaces composed of mainly hydrophobic residues. Interface 1 is the interaction between the two α2 helices from each protein, while Interface 2 involves a four-helix bundle composed of α1 and α4 helices of both proteins [29]. A comparison of the AlphaFold structures of the yeast and *Giardia* orthologues shows that the hydrophobic residues of Interface 1 are largely conserved (Supplementary Fig. 3A). However, there are significant changes at the putative Interface 2 of the *Giardia* protein pair, with several hydrophobic residues being replaced by hydrophilic ones (Supplementary Fig. 3B). Leu13, Leu174 and Leu175 of Rpn8 are occupied by Thr11, Asp174 and Ser175, respectively, in GlRpn8. All these hydrophilic substitutions are located towards the periphery of Interface 2, with the internal hydrophobic residues being conserved, Leu14 in GlRpn8 in place of Leu16 (Fig. 4A), and Val171 instead of Leu171 (data not shown). These changes will likely weaken the strength of the hydrophobic interaction between GlRpn8 and GlRpn11. Besides this hydrophobic patch, the crystal structure further revealed that Interface 2 also involves the insertion of Rpn11’s Met212 into a tight hydrophobic pocket in Rpn8 lined with six residues, Leu15, Leu16, Leu19, Val123, Gln127, and Pro133 [29]. While the methionine residue is conserved in GlRpn11 (Met231) (Supplementary Fig. 4A), Ile13, Leu14, Ala17, Val124, Glu128, and Cys139 line GlRpn8’s hydrophobic pocket (Fig. 4A). The presence of Glu128 will decrease the hydrophobicity of the pocket, further contributing towards the weakening of the interaction between GlRpn11 and GlRpn8. The AlphaFold model of GlRpn8 also indicates that the hydrophobic pocket is less tight than in Rpn8 (Fig. 4B, C). To quantify the size of the hydrophobic pocket opening of GlRpn8 and Rpn8, the minimum pairwise distances of three rim residues were computed across all frames of the last 15 ns of the simulated trajectories. To compare the size of the hydrophobic pocket opening, the normalized probability distributions of all values were estimated for each type of distance and compared between the two systems. Leu16-Pro133 (D1), Leu16-Gln127 (D2), and Gln127-Pro133 (D3) mark the groove size in Rpn8 whereas, the equivalents, Leu14-Cys139 (D1), Leu14-Glu128 (D2), and Glu128-Cys139 (D3) mark the groove size of GlRpn8. All three edges have significantly higher most-probable distance values (∼3-6 Å shifts) in GlRpn8 than Rpn8 (Fig. 4B, C, D). The greater separation of these rim residues suggests that GlRpn8 has a wider hydrophobic pocket. Taking all these differences into account, we conclude that Interface 2-mediated hydrophobic interaction between GlRpn11 and GlRpn8 is likely to be less robust compared to that between the yeast orthologues.

**Fig. 4.**
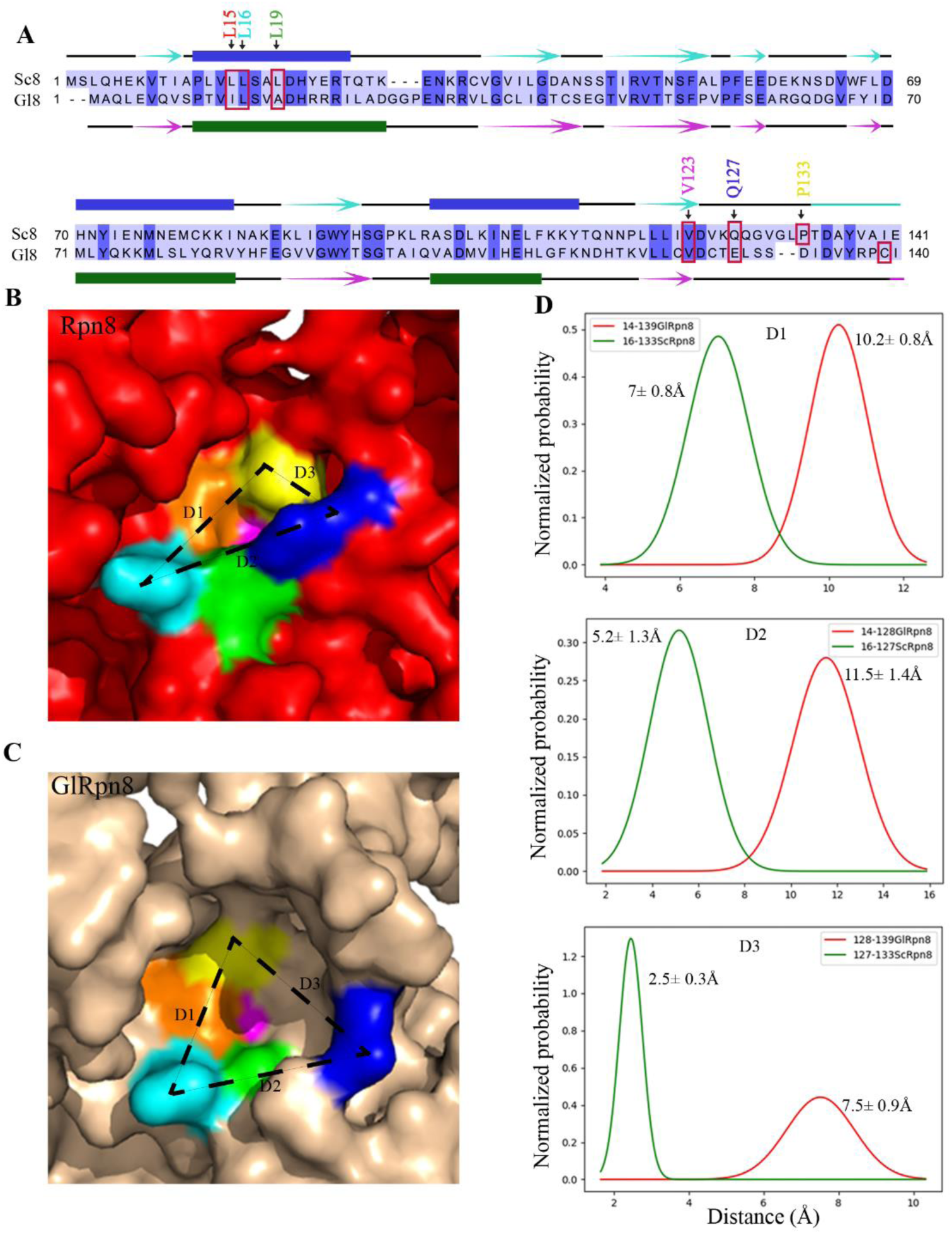
GlRpn8 has a wider hydrophobic pocket than Rpn8. A) Pairwise sequence alignment of a segment of Rpn8 (Sc8) and GlRpn8 (Gl8). The six residues involved in forming the hydrophobic pocket of Rpn8 are boxed in red, and their positions in the protein are indicated above. The predictions of secondary structure elements of each protein, derived from AlphaFold, are given either above (for Sc8) or below (for Gl8), where filled boxes represent α-helices, arrows represent β-strands, and black lines represent unstructured regions. B) and C) Surface representation of the hydrophobic pocket of AlphaFold-derived structures of Rpn8 and GlRpn8. Colour coding of the residues corresponds to that indicated in (A). The dashed lines represent distances between the three residues located along the rim of the hydrophobic pocket. D) Normalized probability distribution of the comparative inter-residue distances from Rpn8 and GlRpn8. Minimum distance measurements between Leu16 and Pro133 (D1), Leu16 and Gln127 (D2), and Gln127 and Pro133 (D3) of Rpn8, derived from its molecular dynamics simulation for 15 ns, were plotted and compared with the corresponding distances values derived from measurements between the residues in GlRpn8, obtained similarly. The raw data underlying the distance plots are provided in Supplementary Data 4.

### 3.4. Extra-proteasomal distribution of GlRpn11

Since the interaction between Rpn8 and Rpn11 is crucial from both the structural and functional properties of the proteasome, the weaker interaction between the giardial orthologues is likely to be of functional significance. Previously, we reported that another subunit of 19S RP, GlRpn10, may have extra-proteasomal functions as extra proteasomal pools of this protein are present at the flagellar pores where the CP subunit Glα7 was not observed [14,53]. We hypothesize that, like GlRpn10, the weak interaction between GlRpn8 and GlRpn11 may have evolved to support an extra proteasomal function. It may be noted that extra proteasomal function has already been documented for Rpn11, which is involved in the fission of mitochondria and peroxisomes, and this activity is independent of its MPN domain-dependent catalytic function [54].

To check if GlRpn11 also has an extraproteasomal distribution, we immunolocalized it with anti-GlRpn11 antiserum. The antiserum detected only a single band of ∼36 kDa in the trophozoite lysate, consistent with the predicted size of GlRpn11 (334 amino acids) (Supplementary Fig. 5). No such band was observed with the pre-immune sera.

Immunolocalization of GlRpn11 in trophozoites showed a diffused nuclear and cytoplasmic distribution, which mirrors a similar distribution of the two other giardial proteasomal components, Glα7 and GlRpn10 and also that of proteasomes from other eukaryotic organisms (Fig. 5A) [14,53]. In addition to the expected nuclear and cytoplasmic distribution, GlRpn11 was present in cytoplasmic puncta, at the V_FP_ and at the OZ of the VD. Given that the Glα7 subunit of the CP is not present at any of these locations, we conclude that these regions contain extra proteasomal pools of GlRpn11. While GlRpn10 is present at all of the four different flagellar pores, GlRpn11 localizes to only the V_FP_ of trophozoites [14]. Additionally, while the flagellar pore signal for GlRpn10 was depleted progressively during the course of encystation, that of GlRpn11 persisted at the V_FP_ and the OZ (Supplementary Fig. 6). This divergent distribution pattern of GlRpn10 and GlRpn11 indicates that both these proteins are likely to discharge non-overlapping extra proteasomal roles.

**Fig. 5.**
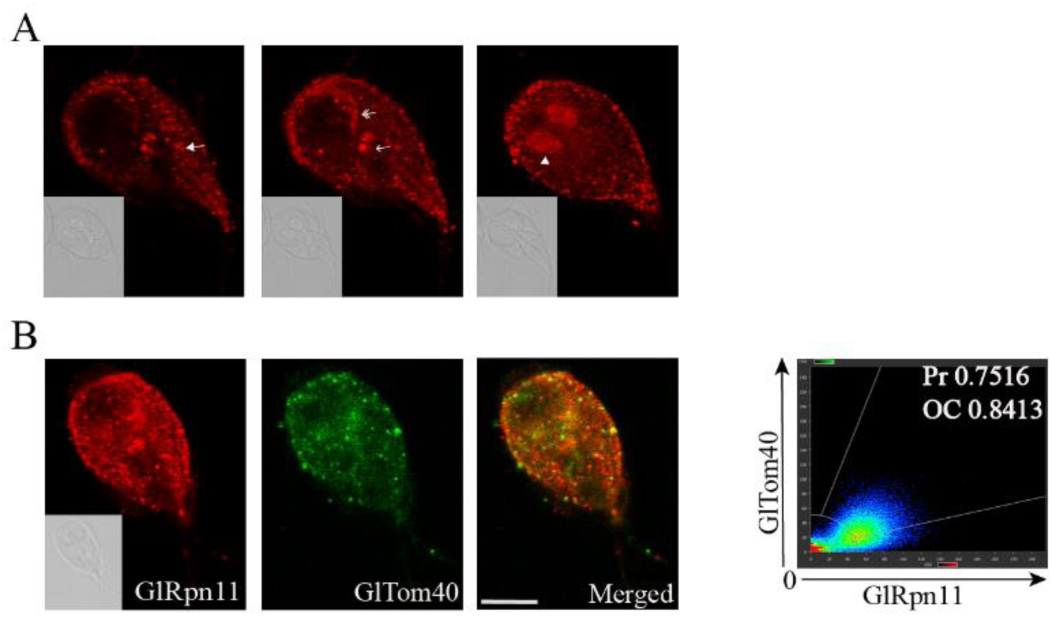
Subcellular distribution of GlRpn11 in trophozoites and its colocalization with GlTom40. A) Immunofluorescence localization of GlRpn11 in trophozoites using polyclonal anti-GlRpn11 antibody. The three panels represent three separate z-sections of an individual image. The solid arrowhead marks the cytoplasmic puncta, the single-headed arrow marks the ventral flagellar pores, the double-headed arrow marks the overlapping zone of the ventral disc, and the triangle marks the nucleus. B) Colocalization of GlRpn11 and GlTom40 in trophozoites. Alexa Flour 594 conjugated anti-rabbit was used to localize GlRpn11 and Alexa Flour 488 conjugated anti-rat was used for GlTom40. The scattergram indicates the colocalization analysis between the two fluorophores in the image, considering all pixels of the cell image. The values for Pearson correlation coefficient (Pr) and overlap coefficient (OC) are indicated. The DIC image of the corresponding z-section is shown in the inset. The scale bar indicates 5 µm.

Studies in yeast show an extra proteasomal function of Rpn11 wherein its C-terminal thirty-one amino acid residues are essential for maintaining mitochondrial morphology and function [24]. While the punctate distribution of GlRpn11 indicated that this protein might also localize to mitosomes, such a role for GlRpn11 cannot be assumed as the C-terminal tails of the yeast and giardial orthologues share very low sequence similarity (Supplementary Fig. 4B). Also, mitochondria and mitosomes differ significantly in both structure and function. While both organelles are bound by a double membrane, mitochondria form a dynamic branched tubular network that undergoes fission and fusion, but mitosomes are isolated spheroidal structures that are static during interphase [55]. In addition, mitosomes are devoid of DNA and lack enzymes of the TCA cycle and components of the electron transport chain [55]. To determine if the cytoplasmic puncta represents a mitosomal distribution of GlRpn11, we colocalized it with GlTom40, a protein import channel of the outer mitosomal membrane that is evolutionarily conserved [56]. Polyclonal anti-GlTom40 detected a prominent band close to the expected size of 39 kDa (Supplementary Fig. 7). Coimmunolocalization of GlRpn11 and GlTom40 revealed that a significant portion of the GlRpn11-positive puncta were also positive for GlTom40 with a Pearson’s Correlation Coefficient of 0.75 (Fig. 5A). Based on the results of the colocalization experiment, we conclude that GlRpn11 localizes to mitosomes, which, along with its localization to the V_FP_ and the OZ indicates the presence of additional extra-proteasomal pools of GlRpn11.

### 3.5. GlRpn11 retains the ability to regulate mitochondrial function

The presence of GlRpn11 at mitosomes led us to hypothesize that while it shares low sequence similarity with Rpn11 at the C-terminus (Supplementary Fig. 4B), it may perform the same function at the mitosome as the yeast protein does at the mitochondria [24]. Rpn11’s role in the maintenance of the mitochondrial network is independent of its deubiquitinase activity as it is mediated by the last thirty-one amino acids, whose deletion causes mitochondrial dysfunction and a change in mitochondrial distribution pattern from tubular to fragmented [24,52]. These phenotypes are rescued by the expression of the missing thirty-one amino acids *in trans* [24]. Since GlRpn11 can substitute for Rpn11’s proteasomal function (Fig. 3A), we visualized the mitochondrial morphology of the *rpn11Δ* strain expressing GlRpn11 for any alteration in mitochondrial distribution compared to the same strain expressing the yeast protein. Mitotracker Deep Red FM staining revealed very little difference in the mitochondrial distribution pattern of the *rpn11Δ* expressing either Rpn11 or GlRpn11 (Supplementary Fig. 8). This indicated that GlRpn1l may have the ability to function both in the context of the proteasome and the mitochondria in yeast.

Based on the ability of the C-terminal residues of Rpn11 to rescue the mitochondrial phenotypes, we adopted a screening strategy to delineate the region of GlRpn11 that can function in the context of the yeast mitochondria. We regenerated the *rpn11-m1* strain described by Rinaldi et al by deleting the chromosomal *RPN11* in cells expressing a truncated Rpn11 lacking the last thirty-one amino acids (Rpn11_1-275_) [24]. Unlike the reticulate mitochondrial distribution pattern of the wild-type, this strain had fragmented mitochondria (Fig. 6A), which is consistent with the observations of Rinaldi et al [24]. Mitochondrial function was independently assessed by measuring endogenous OCR using an Oxytherm device fitted with a Clark electrode. While the OCR of the wild-type yeast was 17.9 nmol/min/10^8^ cells, the value for *rpn11-m1* was significantly lower, at 1.49 nmol/min/10^8^ cells (Fig. 6B). Consistent with the previous report, the reticulate mitochondrial pattern was restored in *rpn11-m1* transformants expressing the C-terminal half of Rpn11, Sc11_140-306_, with an OCR of 16.99 nmol/min/10^8^ cells; this is similar to that of the wild-type. Expression of the C-terminal half of the *Giardia* protein, Gl11_180-334_, partially rescued the fragmented mitochondria phenotype (Fig. 6A), with an OCR value of 8.02 nmol/min/10^8^ cells (Fig. 6B). This indicates that GlRpn11 can not only function in the context of the yeast proteasome but can also partially substitute for the yeast protein’s function in maintaining mitochondrial morphology and function.

**Fig 6.**
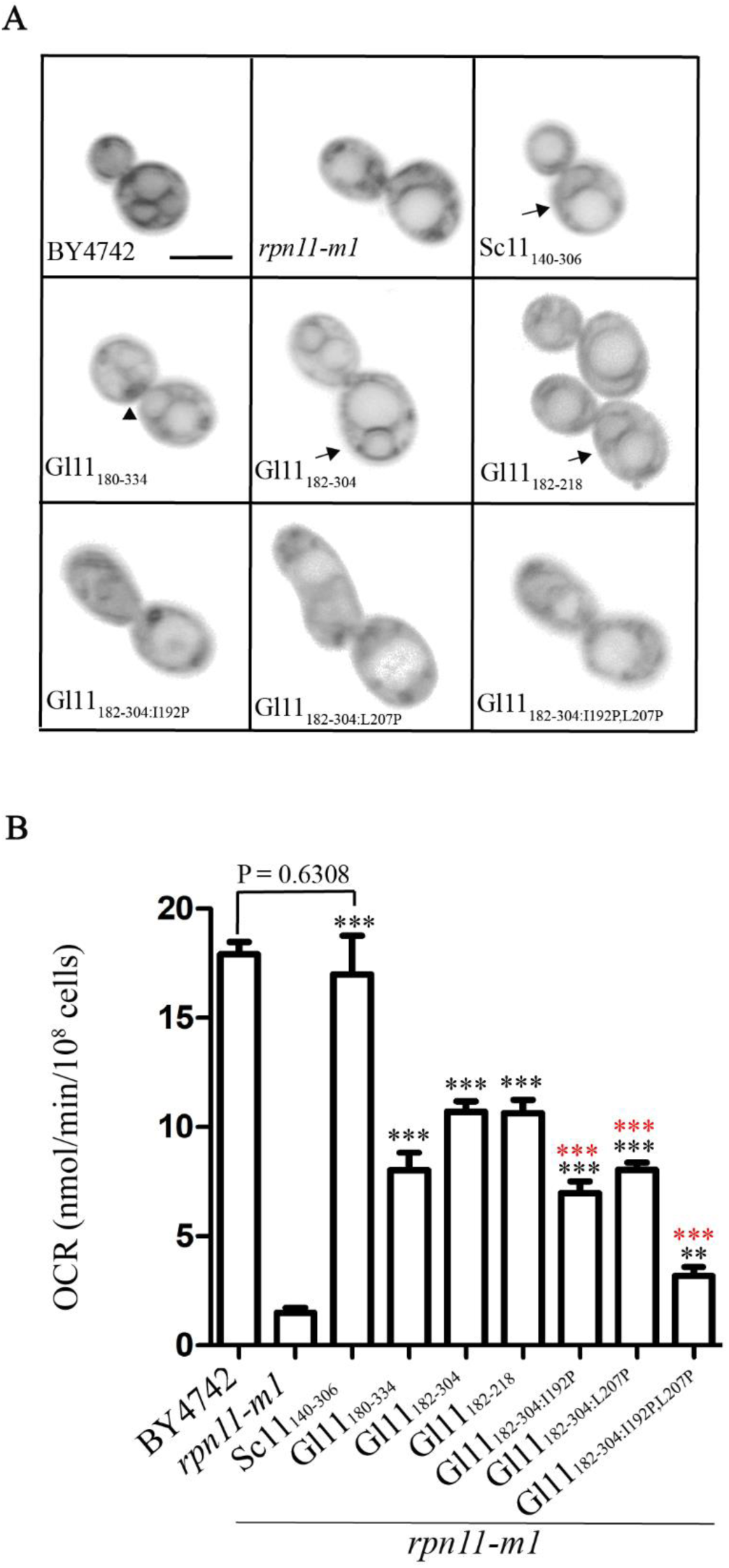
Identification of the region of GlRpn11 that regulates mitochondrial function. A) Monitoring mitochondrial network morphology in cells expressing fragments of Rpn11 and GlRpn11. Mitotracker Deep Red FM was used to monitor the mitochondrial network in wild-type (BY4742) and the *rpn11-m1* mutant that was transformed with a *LEU2* vector expressing various fragments of either Rpn11 (Sc11) or GlRpn11 (Gl11). The fragment length is indicated in the subscript. The bottom three images are of cells that harbour mutant fragments and the positions and amino acid substitutions are also indicated in the subscript. Arrows indicate reticulate mitochondrial morphology, and arrowheads indicate fragmented mitochondria. The scale bar represents 5 µm. B) Measurement of endogenous oxygen consumption rate (OCR) in wild-type and *rpn11-m1* expressing fragments of Rpn11 and GlRpn11. Mean OCR values from three independent observations were plotted, with ±SD. The statistical significance of the difference in OCR is represented as follows: *** *p*-value *<* 0.0001; ** *p*-value *<* 0.001; * *p*-value *<* 0.01; ns *p*-value *>* 0.01, where n = 9. The black asterisk denotes the degree of significance with respect to *rpn11-m1*, and the red asterisk denotes the degree of significance with respect to Glll_182-304._ The raw data underlying this graph are provided in Supplementary Data 5.

Since the last thirty-one amino acids of Rpn11 were essential for maintaining mitochondrial function and morphology [24], we wanted to determine if a comparable stretch of GlRpn11 also harbours a similar function. Curiously, in comparison to Gl11_180-334_, expression of Gl11_182-304_, missing the last thirty-one amino acids, resulted in improved rescue of the mitochondrial phenotype of *rpn11-m1*, both in terms of better restoration of the reticulate mitochondrial distribution and also the elevation of OCR to 10.71 nmol/min/10^8^ cells (Fig. 6A, B). Thus, unlike Rpn11, the segment of GlRpn11 regulating mitochondrial function is not located at its C-terminal end. Further truncations of the protein revealed that Gl11_182-218_ is the smallest fragment that can restore the reticulate mitochondrial morphology (Fig. 6A). This fragment yielded an OCR of 10.64 nmol/min/10^8^ cells, comparable to that observed for Gl11_182-304_ (Fig. 6A). Rinaldi et al documented that an α-helix located near the Rpn11 C-terminus regulated mitochondrial fission [24]. Analysis of the sequence between 182 and 218 of GlRpn11 with Phyre^2^ [25] indicates the possible presence of two short helices (Supplementary Fig. 4A). To determine if these putative helices contribute towards maintaining mitochondrial morphology and function, we substituted a centrally-located residue in each helix of Gl11_182-304_ with a helix disrupter, Pro. Expression of either Gl11_182-304:I192P_ or Gl11_182-304:L207P_ lacked reticulate mitochondrial morphology (Fig. 6A) and their OCR values were lower than the unmutated fragment, being 6.97 nmol/min/10^8^ cells for Gl11_182-304:I192P_ and 8.03 nmol/min/10^8^ cells for Gl11_182-304:L207P_ (Fig. 6B). The double mutant, Gl11_182-304:I192P, L207P_ not only displayed fragmented mitochondria but had a significantly lower OCR of 3.19 nmol/min/10^8^ cells. Thus, the two predicted helices, which together span a large part of the minimal region of GlRpn11 (182-218) that can sustain mitochondrial morphology and function, play a crucial role in this activity as their disruption in even the larger fragment of GlRpn11 (182-304) significantly reduces the ability of the fragment to rescue mitochondrial function. However, since the OCR value of the double mutant is higher than that of *rpn11-m1*, it is possible that other segments of the Gl11_182-304_ may contribute to mitochondrial function. Taken together, the above results indicate that an internal segment of GlRpn11 has the ability to support yeast mitochondrial structure and function. While the position of the sequence capable of such function varies between Rpn11 and GlRpn11, the secondary structure of such region in each respective protein is similar. Given that pools of GlRpn11 are present at the mitosomes and the protein has the ability to modulate the structure and function of yeast mitochondria, we propose that, like Rpn11, GlRpn11 may modulate mitosomal function in *Giardia*. However, the evolutionary divergence between yeast and *Giardia* has resulted in not only a change in the position of the motif/domain within the two orthologues but also how their function is regulated as the C-terminal end of GlRpn11 appears to have a negative regulatory role, whereas no such segment has been documented for Rpn11.

## 4. Discussion

This study was initiated to understand if, in the absence of Rpn12 and Sem1, the interactions within the proteasome lid particle of *Giardia* are different from those present in yeast. Our data indicates that in many cases, while there are interactions between two subunits in yeast and *Giardia*, the affinity between the giardial protein pairs is higher (subunits connected by green lines in Fig. 7). We also observed weak interactions between some giardial subunits, where there was none between the yeast orthologues (blue lines in Fig. 7). In only two cases, we observed weaker interactions between the *Giardia* proteins, compared to the strong interactions between the yeast pair (red lines in Fig. 7). Hence, this comparative analysis of the binary protein interaction affinities between the lid particle components from these two organisms reveals higher affinities between several giardial proteasome lid subunits. These together will likely form a stable lid complex without Rpn12 and Sem1. Incidentally, *RPN12* is an essential gene in yeast [57], and the absence of the orthologous gene in *Giardia* indicates that its requirement may be bypassed by altering the inter-subunit interactions between the other lid subunits.

**Fig 7.**
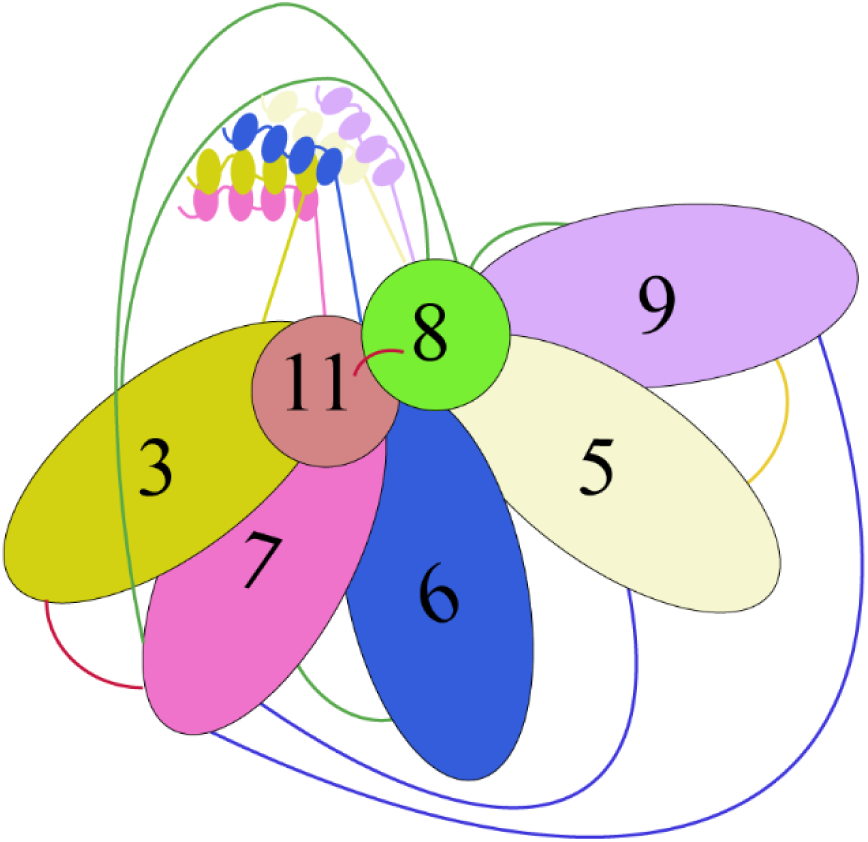
Comparative analysis of binary interaction affinities between the proteasome lid subunits from yeast and *Giardia*. Schematic representation of the interactions detected between the proteasome lid subunits of *Giardia lamblia*. The lines connect subunits where binary interactions have been detected between the *Giardia* proteins. These lines are colour-coded to compare binary affinities between orthologous proteins from yeast and *Giardia*. Green: interactions that are stronger in *Giardia* than in yeast; blue: interactions observed only in *Giardia* but not in yeast; yellow: interaction of equivalent affinity in yeast and *Giardia*; red: interactions that are weaker in *Giardia* than in yeast.

Structural studies of the yeast proteasome lid indicate that Rpn12 and Sem1 occupy peripheral positions [47,58]. Also, these two proteins are smaller than the remaining proteasomal lid subunits, being 274 and 89 amino acid residues in length, respectively. Consistent with their small size and peripheral positions, they make limited contact with other lid components [47]. The PCI domain of Rpn12 has maximum contact with Rpn3, and its C-terminal α-helix contacts the helices of Rpn3, Rpn8, Rpn9 and Rpn11 [47]. Sem1 makes contact with Rpn3 and Rpn7 only [15,47]. Thus, the loss of these two subunits is unlikely to entail major structural changes in the proteasome lid, making it likely that enhanced interactions between the remaining lid components can compensate for the requirement for these subunits.

Of the few interactions that are weaker compared to those in yeast, that between GlRpn11 and GlRpn8 was notable, as the strong interaction between the Rpn8-Rpn11 dimer forms a platform for lid assembly in yeast [59]. The weak interaction between GlRpn8 and GlRpn11 may have evolved to support the extra proteasomal role of the latter at the OZ of the VD, the V_FP_, and the mitosomes. However, this weak interaction between GlRpn8 and GlRpn11 does not appear to hamper their ability to function in the context of the yeast proteasome, as each can functionally complement the corresponding deletion mutant (Fig. 3). In addition, GlRpn11 also discharges the extra-proteasomal functions of Rpn11. While Rpn11 also performs an extra proteasomal function at the mitochondria, such functions of GlRpn11 appear to be more extensive as, besides the mitosomes, substantial pools of this protein are also present at the OZ and the V_FP_. The weak interaction between GlRpn8 and GlRpn11 may have evolved to increase the proteasome-independent pool of GlRpn11. It may be noted that Alvarado et al have reported the presence of GlRpn11 at the ventral flagella and nuclear foci of trophozoites, which we have not observed [18]. Nor have we observed the oscillating presence of GlRpn11 at various flagellar locations during the course of encystation [18]. Instead, similar to the previously documented distribution of Glα7 [53], we have observed that GlRpn11 also localizes to larger encystation-specific vesicle-like structures after induction of encystation (Supplementary Fig. 6).

The absence of Sem1 in *Giardia* may also augment such proteasome-independent pools. Studies in *Aspergillus nidulans* have shown that Sem1 is required for stable interaction of Rpn11 and Rpn10 with the 19S proteasome lid [60]. As previously mentioned, extra-proteasomal pools of GlRpn10 are present at all the flagellar pores [14]. However, other than GlRpn10, no extra proteasomal function has been documented for any Rpn10 orthologue. The absence of Sem1 in *Giardia* likely allows both GlRpn11 and GlRpn10 to dissociate from the proteasome and perform extra proteasomal functions unique to this organism. Given the presence of GlRpn10 and GlRpn11 at locations that are specific to *Giardia*’s cell shape (except for location at the mitosomes), the absence of Sem1 may, in fact, contribute to the unique cellular architecture of this organism. Whether reductive evolution resulted in the loss of Rpn12 and Sem1 from the genome of *Giardia*’s ancestor, remains to be determined. Thus, our study has not only uncovered how the interactions within the proteasome lid of *Giardia* have altered in the absence of Rpn12 and Sem1 but also revealed how GlRpn11 may dissociate from the proteasome to function at specific subcellular locations.

Given the functional association of Rpn11 with mitochondria [24], the mitosomal location of GlRpn11 indicates that the role of this protein in mitosomal fission has been retained even after the reductive evolution of *Giardia* during which mitosomes, which are remnants of mitochondria, themselves have lost their entire genome and a sizable part of their proteome [55]. It may be noted that while functional complementation shows that both the Rpn11 orthologues from yeast and *Giardia* can sustain yeast mitochondrial function, the extreme C-terminal end of GlRpn11 may play a negative regulatory role in this process (Fig. 6). This may represent a novel regulatory circuit that allows *Giardia* to control mitosomal fission.

The presence of extra proteasomal pools of GlRpn11 indicates that it has sequences that support its targeting to the mitosome, the OZ and the V_FP_. The OZ is part of the VD, which is unique to *Giardia* as this appendage is absent in even the closely related *Salmonicida* species belonging to the same Hexamitidae family [61]. The lack of detection of this protein in previous proteomic analyses of the isolated VD indicates that its association with the structure is not stable [62]. Further studies are needed to reveal the role of GlRpn11 at these extra proteasomal locations. Also, identifying the regions of GlRpn11 that play a role in its distribution to multiple subcellular locations and the other cellular factors that assist in this process will shed light on the underlying regulatory process.

Our functional complementation studies showing that a fragment of GlRpn11 sustains the yeast mitochondrial morphology and function indicates that the role of GlRpn11 at the mitosome is likely independent of its deubiquitinase activity. Whether it functions as a proteasome-independent deubiquitinase, the OZ, or the V_FP_ remains to be determined. However, based on the limited role of ubiquitin machinery in regulating the cellular functions of *Giardia*, we speculate that GlRpn11’s functional role at these locations is unlikely to involve its deubiquitinase function. The extra-proteasomal functions of GlRpn11 indicate that it is a moonlighting protein. Our previous report documenting the extra proteasomal localization of GlRpn10 shows that it, too, may be capable of functions beyond serving as a ubiquitin receptor [14]. Such moonlighting functions of certain proteins, in addition to the well-established canonical functions gleaned from studies in model organisms, may enable this unicellular eukaryote to sustain the unique cellular features that enable it to sustain its parasitic lifecycle wherein it must survive within the host gut and also maintain two morphological states. The size of the reference genome of the WB isolate of *Giardia* is 11.7 Mb, which is ∼250X smaller than the human genome [63]. Yet this small genome sustains not only the complex cellular morphology and motility of trophozoites but also its transition to cysts that entails the retraction of the flagella, the disassembly of the VD and the change in the shape of the cell [64]. The morphology of the trophozoite is particularly intriguing as this unicellular eukaryote has a bilaterally symmetrical ‘inverted tear-drop’ cell shape, where even the four different flagella pairs emerge symmetrically from the cell. In addition, each flagella pair displays unique movement that allows the parasite to display a range of movements [65]. All these complex cellular features and functions are made possible even when the genome encodes a limited set of proteins. Since most *Giardia* genes lack introns, alternative splicing cannot be used to make alternative forms of proteins from the same genome segment [63]. Thus, moonlighting functions of proteins may be a strategy that allows *Giardia* to successfully thrive in the niche environment of the host gut and spread from host to host.

The fundamental cellular function of the proteasome is the reason for its ubiquitous presence in all eukaryotes. This study highlights that the architecture of such ubiquitous protein complexes need not be the same in all organisms. Variations of both the CP and RP of the proteasome have already been documented in mammalian systems [66]. Even though there is extensive literature documenting the architecture of these different proteasomes from mammalian model organisms and the unicellular yeast, which is also part of the same opisthokonta supergroup as mammals, very little is known about the subunit compositions of proteasomes from other eukaryotic supergroups. This is the first study to document that it may be possible to assemble a proteasome even without Rpn12, an essential proteasome component in the widely studied model eukaryotes. It highlights the utility of performing comparative studies of fundamental processes in organisms distantly related to established model eukaryotes. Such studies will not only help us understand the different architectures of multisubunit complexes but also pave the way to differentiate between subunits that are essential and those that are not.

## Funding

This work was funded by the Department of Science and Technology (DST-SERB) [CRG/2018/003030/HS] and Bose Institute, Kolkata. AD was supported by the INSPIRE program from the Department of Science and Technology (DST), Government of India [IF170741]. AR [JAN2012-353894] and SM [MAY2018-353734] are supported by the University Grants Commission (UGC). NC is supported by the Council of Scientific & Industrial Research (CSIR) [09/015(0546)/2019].

## Authors’ Contributions

Ankita Das: Conceptualization, Data curation, Formal analysis, Investigation, Methodology, Visualization, Writing - Original draft

Atrayee Ray: Conceptualization, Data curation, Investigation, Visualization

Nibedita Ray Chaudhuri: Data curation, Formal analysis, Investigation, Methodology, Visualization, Writing – original draft

Soumyajit Mukherjee: Data curation, Formal analysis, Methodology, Shubhra Ghosh Dastidar: Formal analysis, Investigation

Alok Ghosh: Investigation

Sandipan Ganguly: Methodology, Resources

Kuladip Jana: Methodology, Investigation

Srimonti Sarkar: Conceptualization, Formal analysis, Project administration, Supervision, Investigation, Resources, Writing - original draft preparation, Writing - review and editing

## Acknowledgments

We thank Prof Alok Kumar Sil for his valuable comments during the course of the study and for his critical comments on the manuscript. DNA sequencing and confocal imaging were conducted in the Central Instrumentation Facility of Bose Institute. We thank Prantik Saha and Sheolee Ghosh-Chakraborty for confocal imaging and Leica Microsystems for technical assistance during image processing. Antibody generation was performed at the Centre for Translational Animal Research, Bose Institute. We thank the members of the Sarkar Laboratory for providing valuable comments during the course of the study.

## Supplementary material

**Supplementary Table 1:** Comparison (percentage of identity) of the 19S lid subunits from *S. cerevisiae, G. lamblia, H. sapiens*.

| Subunit Nomenclature | ORF ID |  |  | Identity score in percentage (local alignment) |  |
| --- | --- | --- | --- | --- | --- |
|  | *S. cerevisiae* | *G. lamblia* | *H. sapiens* | *S. cerevisiae* with *G. lamblia* | *S. cerevisiae* with *H. sapiens* |
| **19S RP base components** |  |  |  |  |  |
| Rpt1 | YKL145 W | GL50803_86683 | NM_002803.4 | 43.9 | 70.0 |
| Rpt2 | YDL007 W | GL50803_113554 | NM_002802.3 | 46.4 | 71.4 |
| Rpt3 | YDR394 W | GL50803_7950 | NM_006503.4 | 44.8 | 66.4 |
| Rpt4 | YOR259C | GL50803_21331 | NM_002806.3 | 42.4 | 60.9 |
| Rpt5 | YOR117<br>W | GL50803_4365 | NM_002804.5 | 40.6 | 65.6 |
| Rpt6 | YGL048C | GL50803_17106 | NM_002805.6 | 53.3 | 72.3 |
| Rpn1 | YHR027C | GL50803_33166 | NM_002808.5 | 19.7 | 21.3 |
| Rpn2 | YIL075C | GL50803_91643 | NM_00119103<br>7.2 | 18.4 | 19.3 |
| Rpn13 | YLR421C | - | NM_175573.2 | - | 27.4 |
| **19S RP Lid components** |  |  |  |  |  |
| Rpn3 | YER021<br>W | GL50803_15454 | NM_002809.4 | 19.8 | 33.9 |
| Rpn5 | YDL147<br>W | GL50803_16929 | NM_002816.5 | 18.9 | 41.5 |
| Rpn6 | YDL097C | GL50803_16659 | NM_00127048<br>2.2 | 19.5 | 43.3 |
| Rpn7 | YPR108W | GL50803_4331 | NM_00127177<br>9.2 | 20.6 | 39 |
| Rpn8 | YOR261C | GL50803_7896 | NM_002811.5 | 28 | 49.3 |
| Rpn9 | YDR427<br>W | GL50803_9099 | NM_002817.4 | 18.5 | 31.8 |
| Rpn10 | YHR200<br>W | GL50803_15604 | NM_00133069<br>2.2 | 22.6 | 51.56 |
| Rpn11 | YFR004W | GL50803_16823 | NM_005805.6 | 40.4 | 66.2 |
| Rpn12 | YFR052W | - | NM_002812.5 | - | 32.1 |
| Rpn15 | YDR363<br>W-A | - | NM_005047.4 | - | 34.1 |

**Supplementary Table 2.**
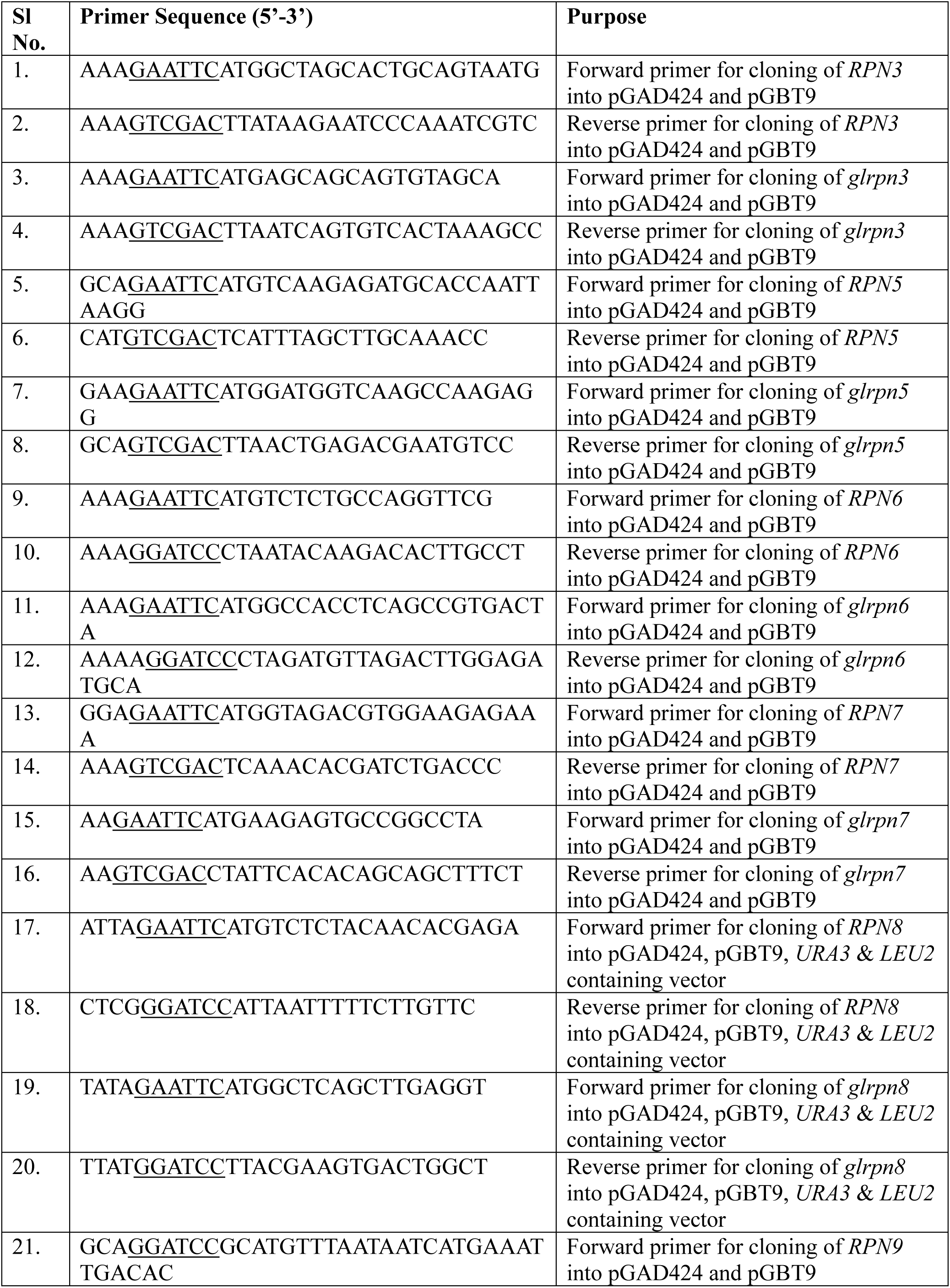
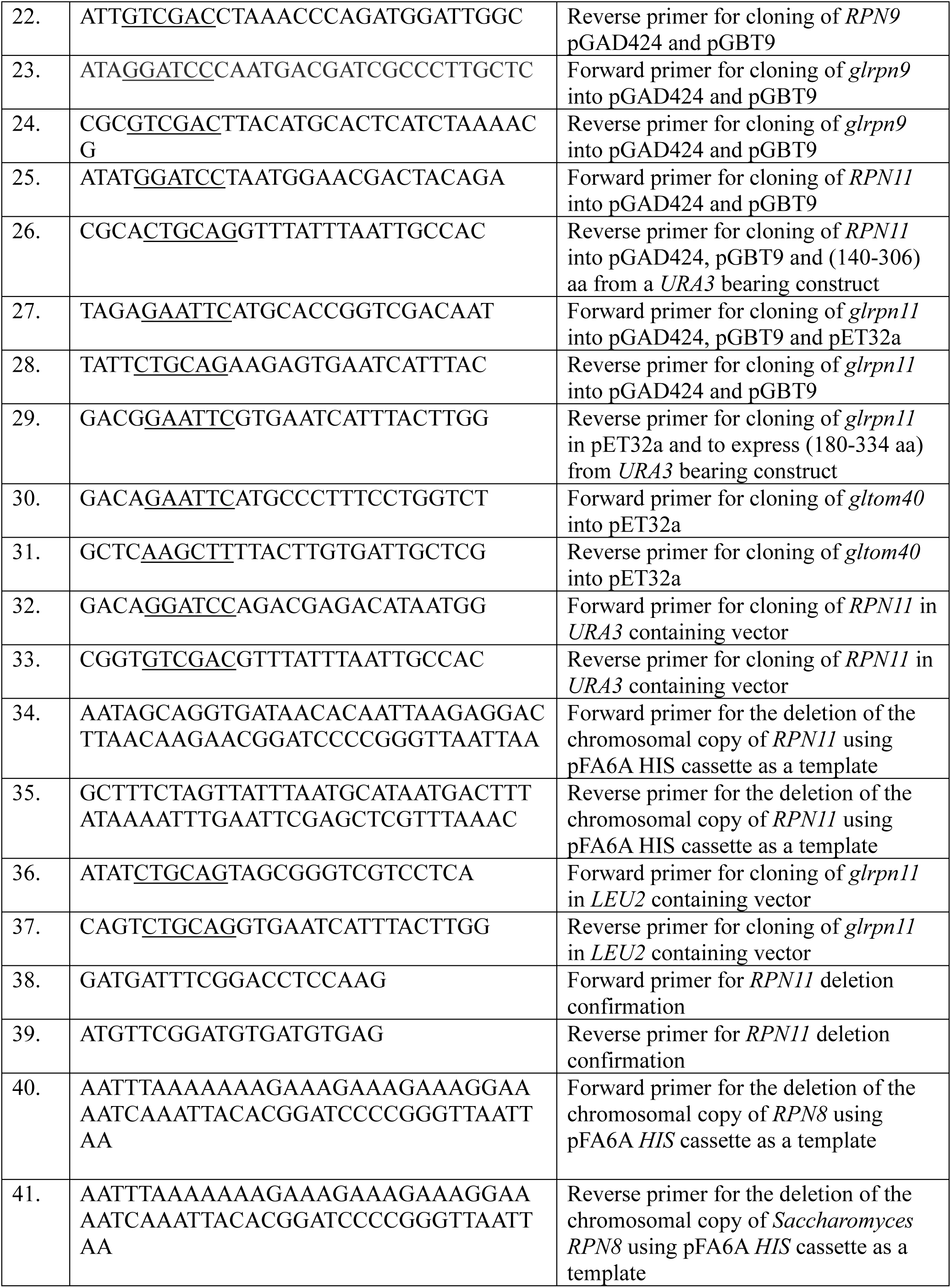
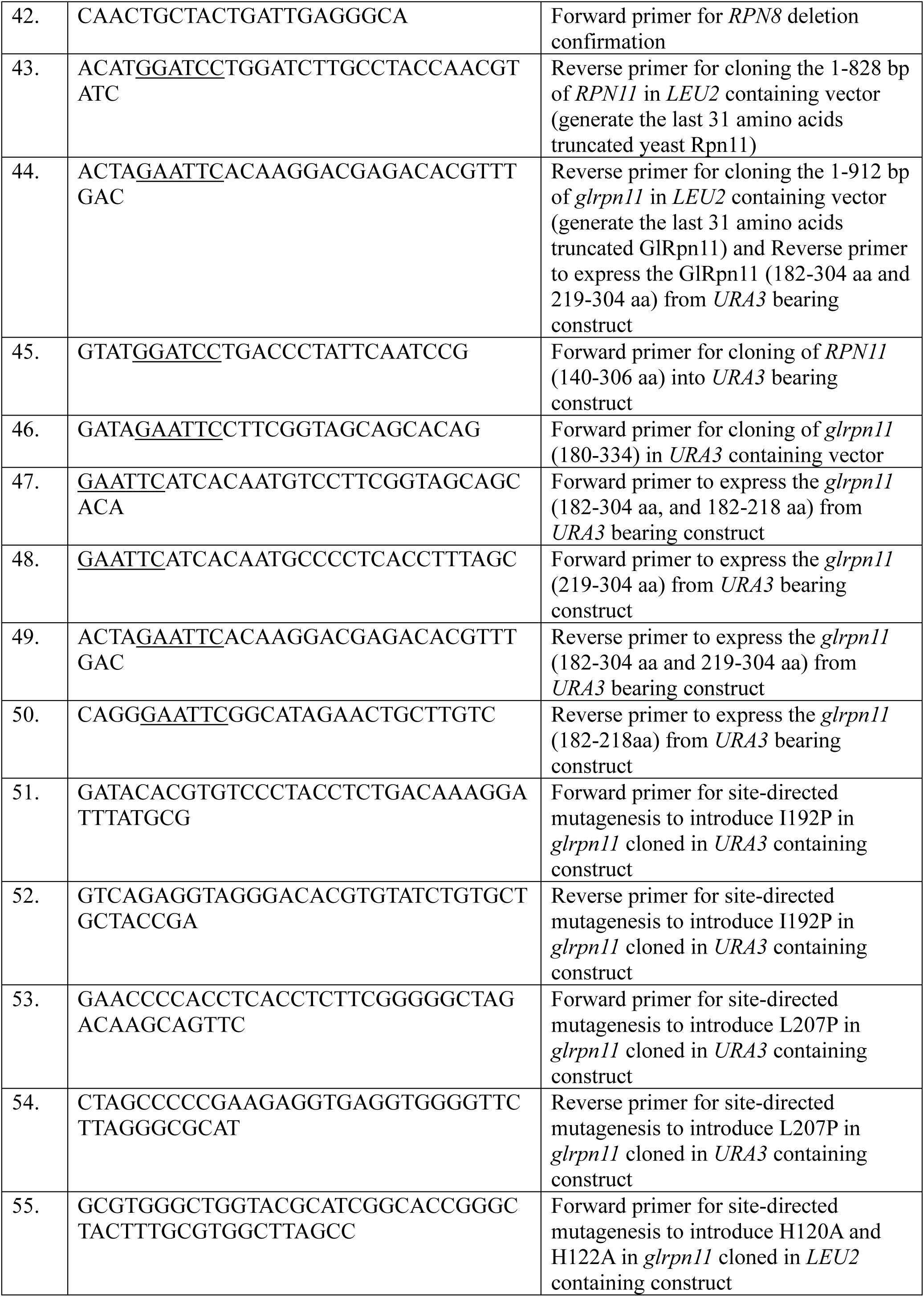
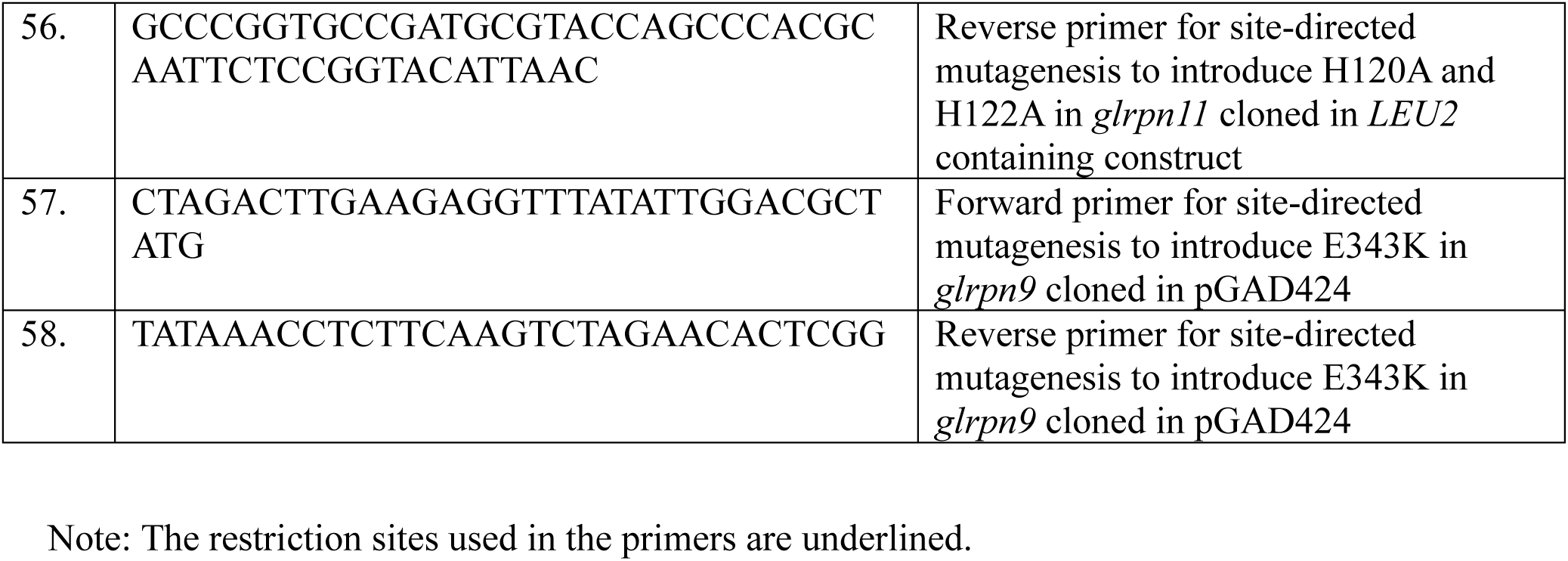
Primer sequences used in this study.

**Supplementary Table 3:**
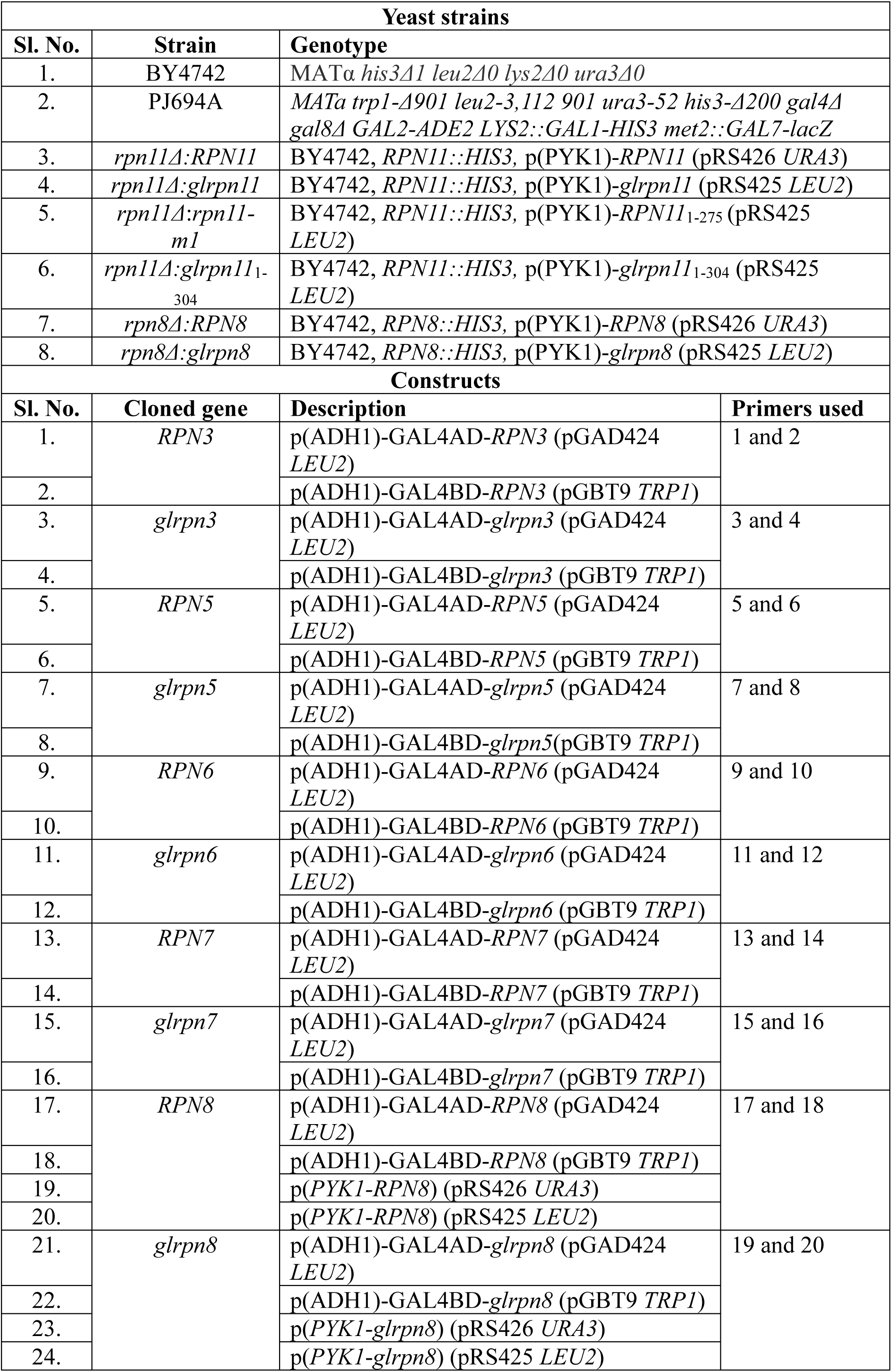
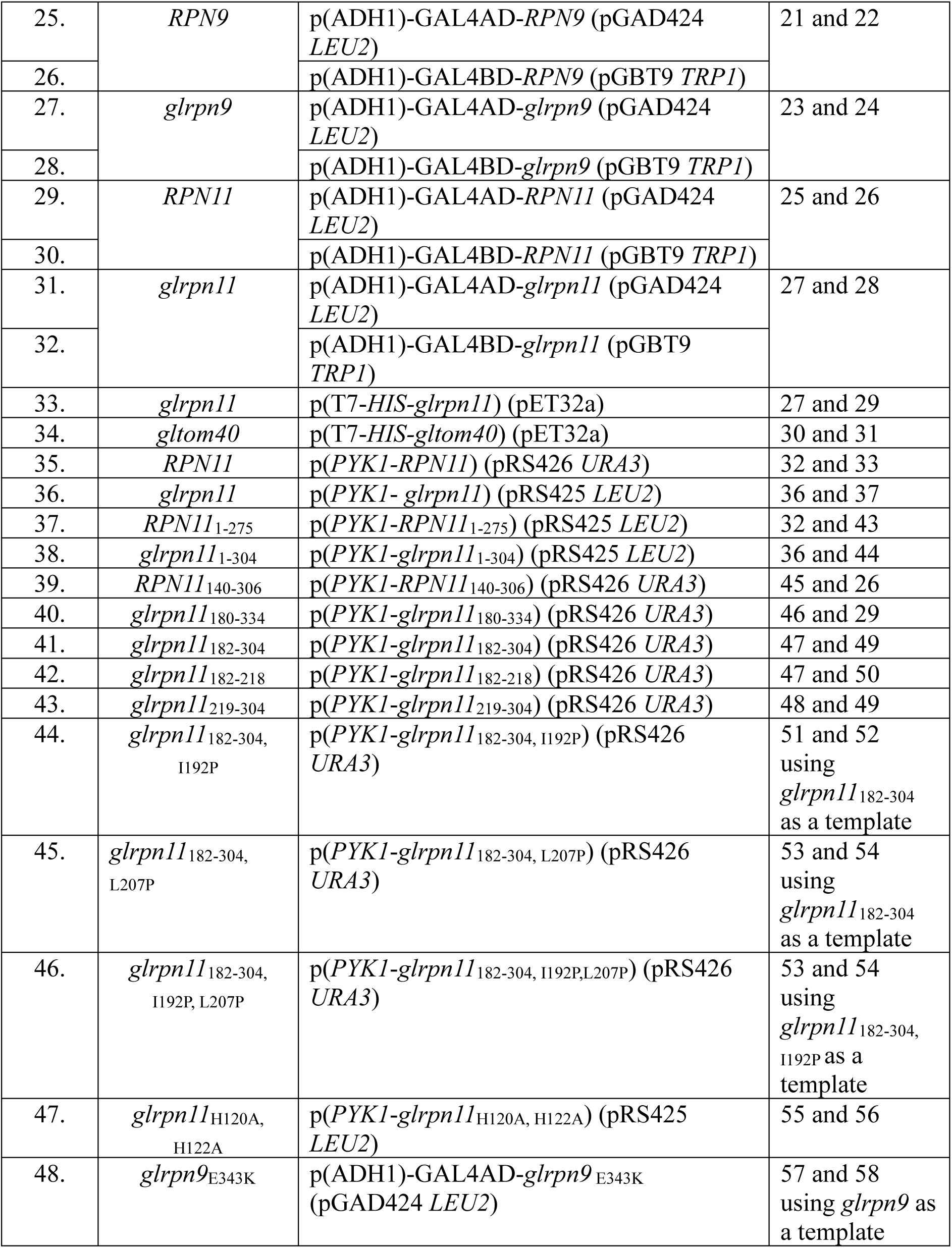
List of yeast strains and constructs used in this study.

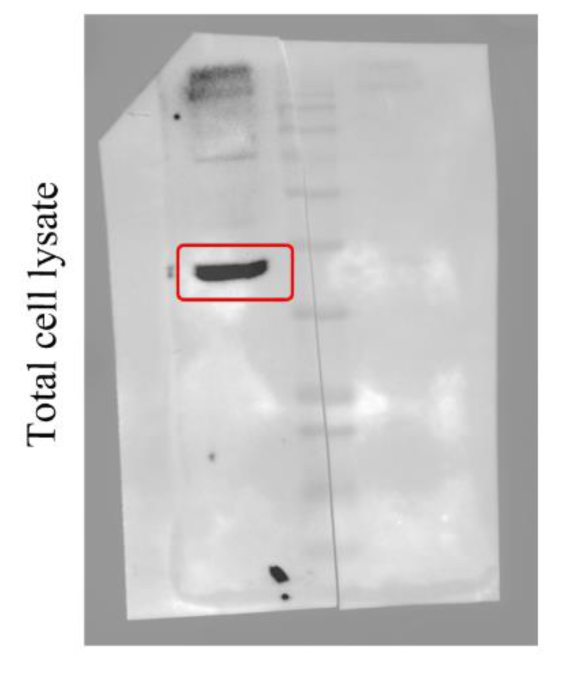

GlRpn11 detection from total cell lysate of *Giardia* trophozoite from Supplementary Fig. 5

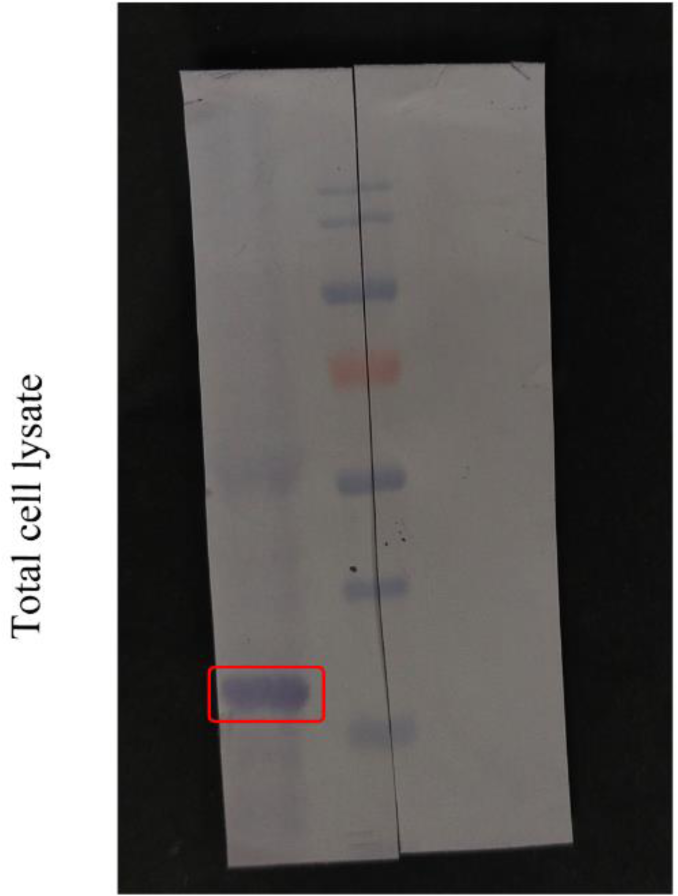

GlTom40 detection from total cell lysate of *Giardia* trophozoite from Supplementary Fig. 7

**Supplementary Data 1: Raw uncropped western blot images**

**Supplementary Data 2. Raw data associated with Fig. 1 (.PZF)**

**Supplementary Data 3. Raw data associated with Fig.2 (PZF)**

**Supplementary Data 4. Raw data associated with Fig. 4 (XLSX)**

**Supplementary Data 5. Raw data associated with Fig. 6 (PZF)**

## Supplementary Figures

**Supplementary Figure 1.**
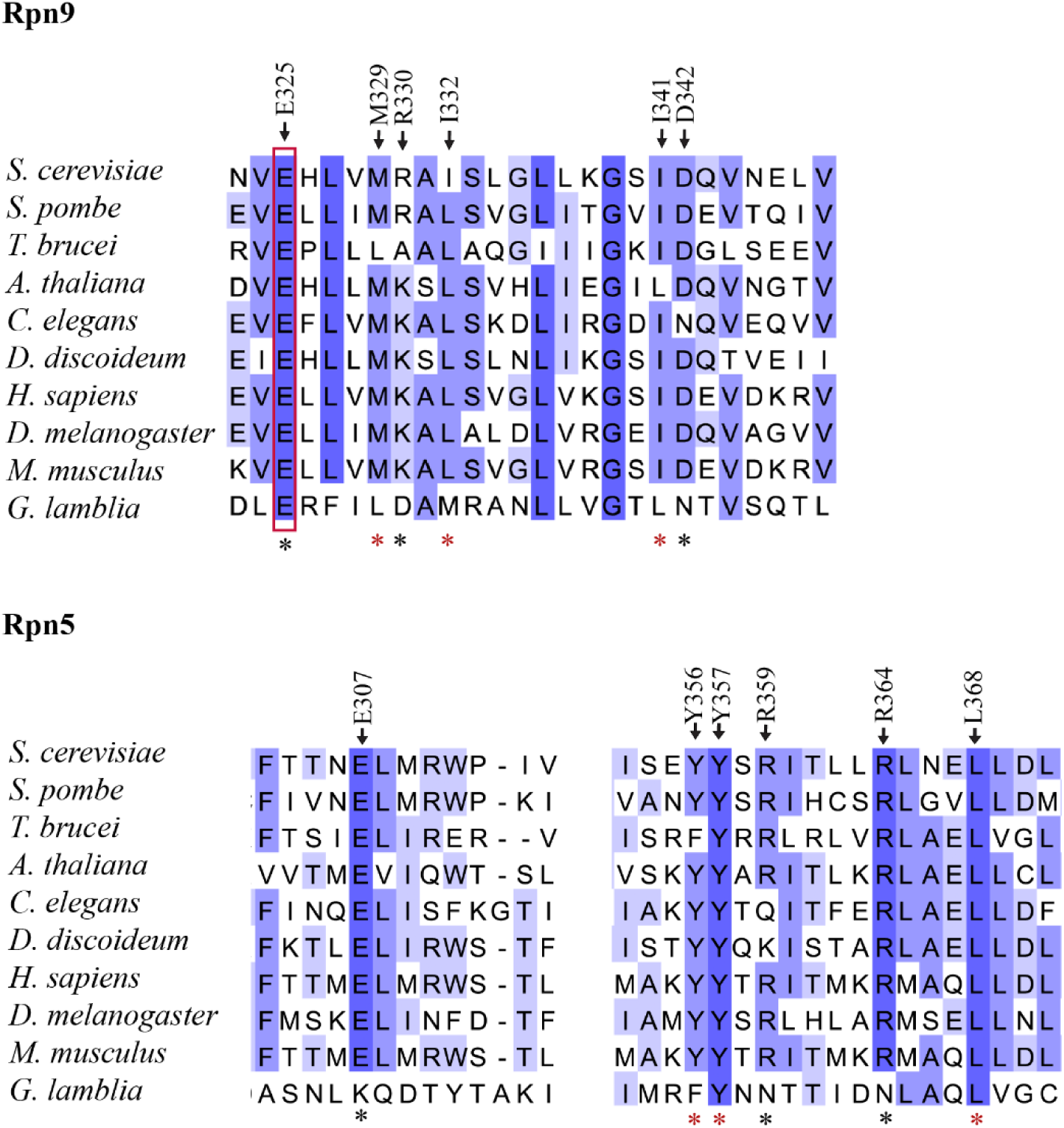
Multiple sequence alignment of fragments from Rpn9 and Rpn5 orthologues. Multiple sequence alignment (MSA) of Rpn9 and Rpn5 orthologues from different taxonomic groups (*Saccharomyces cerevisiae, Schizosaccharomyces pombe, Trypanosoma brucei, Arabidopsis thaliana, Caenorhabditis elegans, Dictyostelium discoideum, Homo sapiens, Drosophila melanogaster, Mus musculus,* and *Giardia lamblia*). The MSA of Rpn9 spans the fragment of *S. cerevisiae* Rpn9 that interacts with Rpn5, while the MSA of Rpn5 spans the fragment of *S. cerevisiae* Rpn5 that interacts with Rpn9. Black asterisks indicate the polar residues, and red asterisk indicates nonpolar residues. The conserved Glu residue mutated to generate fusion protein AD-GlRpn9* (Glu343Lys) is boxed in red. Position identifiers of residues are in accordance with those in *S. cerevisiae* orthologues.

**Supplementary Figure 2.**
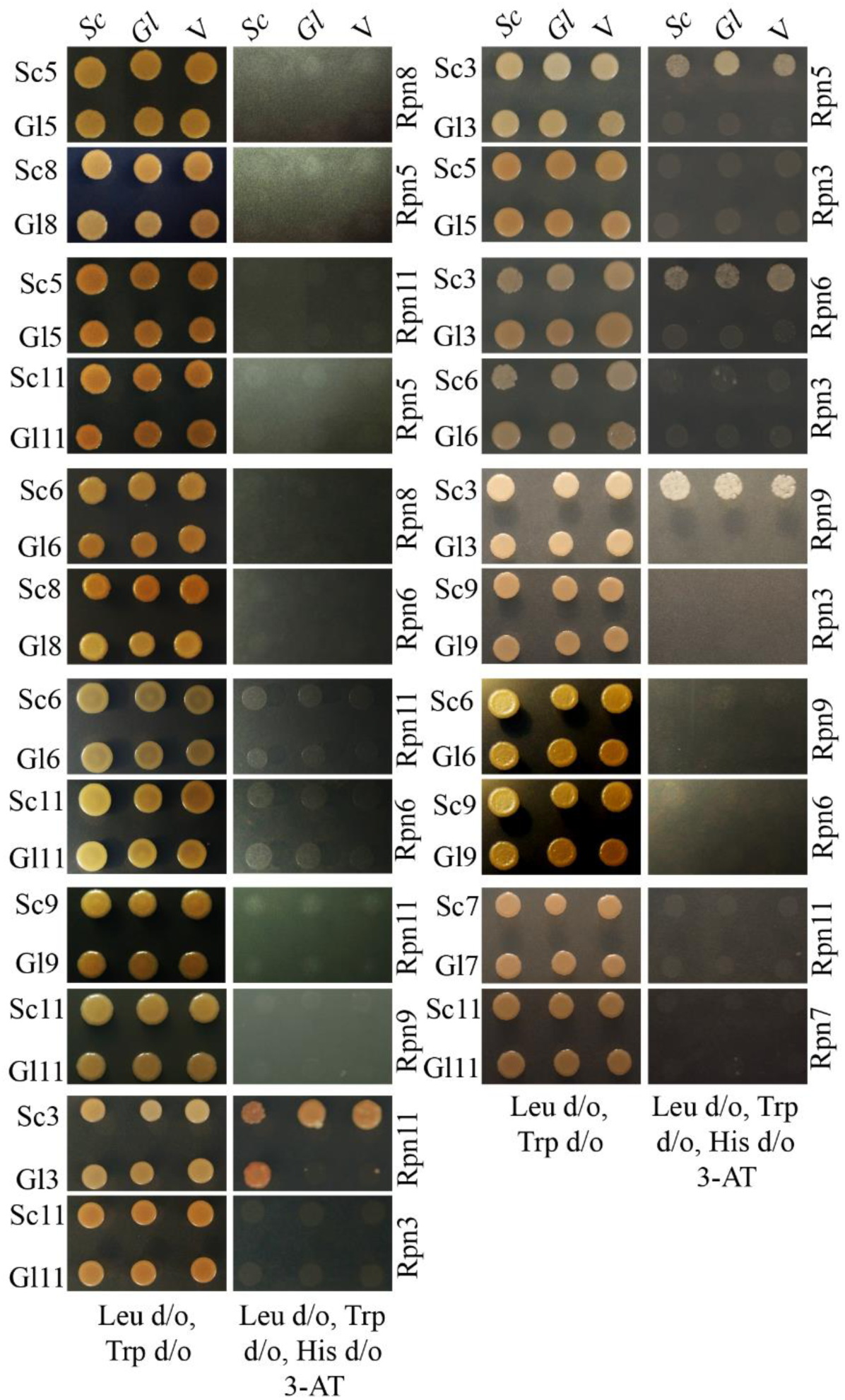
Proteasome lid subunit pairs not showing interaction in yeast two-hybrid assay. The interactions between various proteasome lid subunits of *S. cerevisiae* (Sc) and *G. lamblia* (Gl) were assessed as described in Fig. 1(**A**), 2(**A**). Proteins with the BD domain fused to their N-terminus are indicated on the left of each panel (Sc5, Gl5, etc.), while proteins with the AD domain fused to their N-terminus are indicated to the right of each panel (Rpn5, Rpn8, etc). Growth on media lacking leucine and tryptophan (Leu d/o, Trp d/o) indicates the presence of the AD and BD constructs. Absence of growth on 3-amino-1,2,4-triazole containing media lacking leucine, tryptophan and histidine (Leu d/o, Trp d/o, His d/o 3-AT) indicates the lack of expression of the *HIS3* reporter gene. All plates were incubated at 28°C for 3 days. (V) dictates empty vector pGAD424.

**Supplementary Figure 3.**
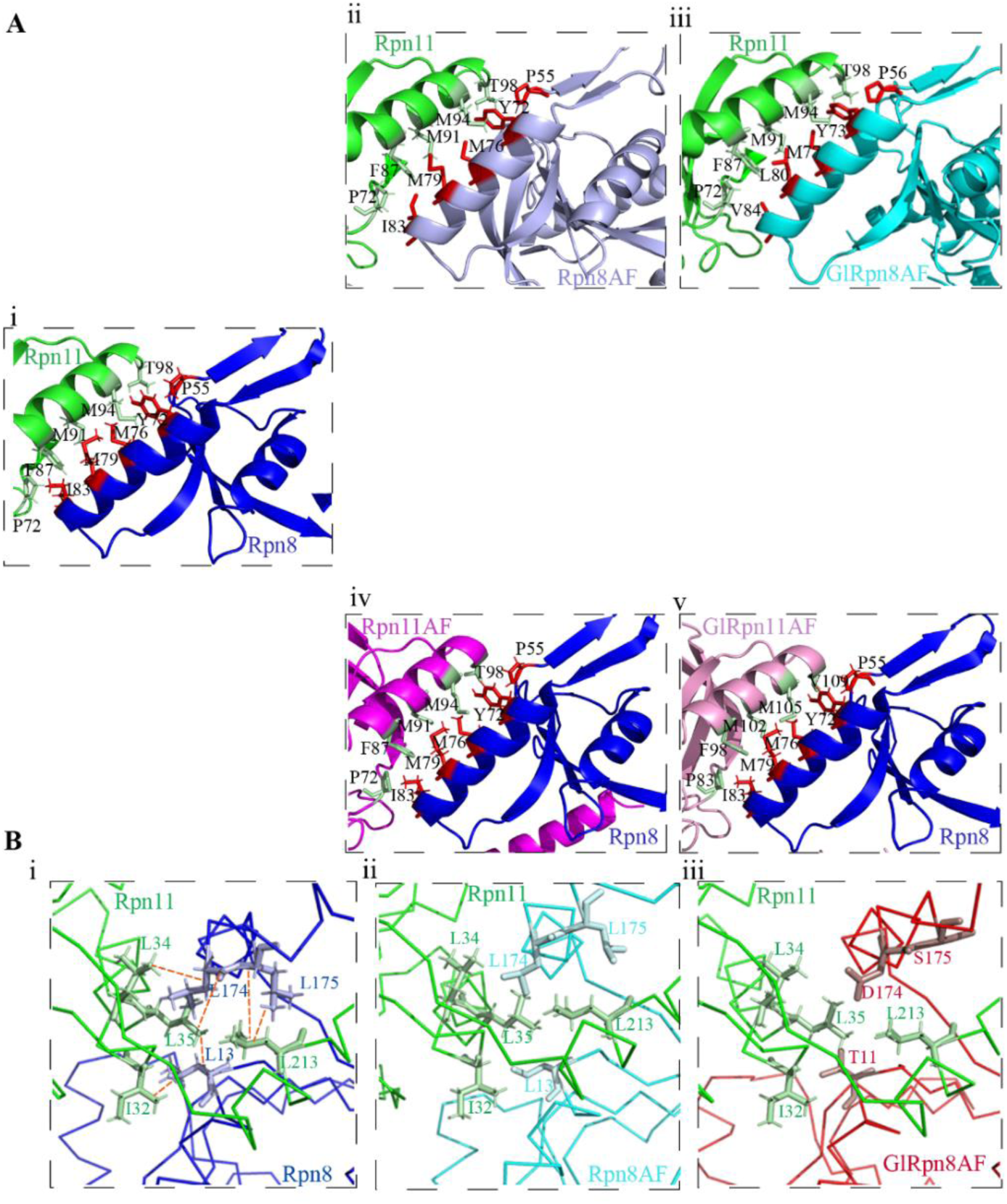
Comparison of Interface 1 and Interface 2 of the predicted interaction surface between GlRpn8 and GlRpn11 with those of Rpn8-Rpn11. **Ai**) Cartoon representations of Interface 1 between Rpn8 and Rpn11 (PDB ID: 4O8X) where Rpn11 backbone is green and Rpn8 backbone is blue. The residues from each protein contributing to the interaction are shown in stick representation. **Aii** and **Aiii**) The crystal structure of Rpn8 (4O8X) in (**Ai**) was replaced with the AlphaFold-predicted structures of either Rpn8 (Rpn8AF) or GlRpn8 (GlRpn8AF). (**Aiv** and **Av**). The crystal structure of Rpn11 (4O8X) in (i) was replaced with the AlphaFold-predicted structures of either Rpn11 (Rpn11AF) or GlRpn11 (GlRpn11AF). (**Bi**) Ribbon representations of Interface 2 between Rpn8 and Rpn11 (PDB ID: 4O8X) where Rpn11 backbone is green and Rpn8 backbone is blue. The residues from each protein contributing to the interaction are shown in stick representation. Hydrophobic interactions are marked with orange dashed lines. (**Bii** and **Biii**) The crystal structure of Rpn8 (4O8X) in (**Bi**) was replaced with the AlphaFold-predicted structures of either Rpn8 (Rpn8AF) or GlRpn8 (GlRpn8AF)

**Supplementary Figure 4.**
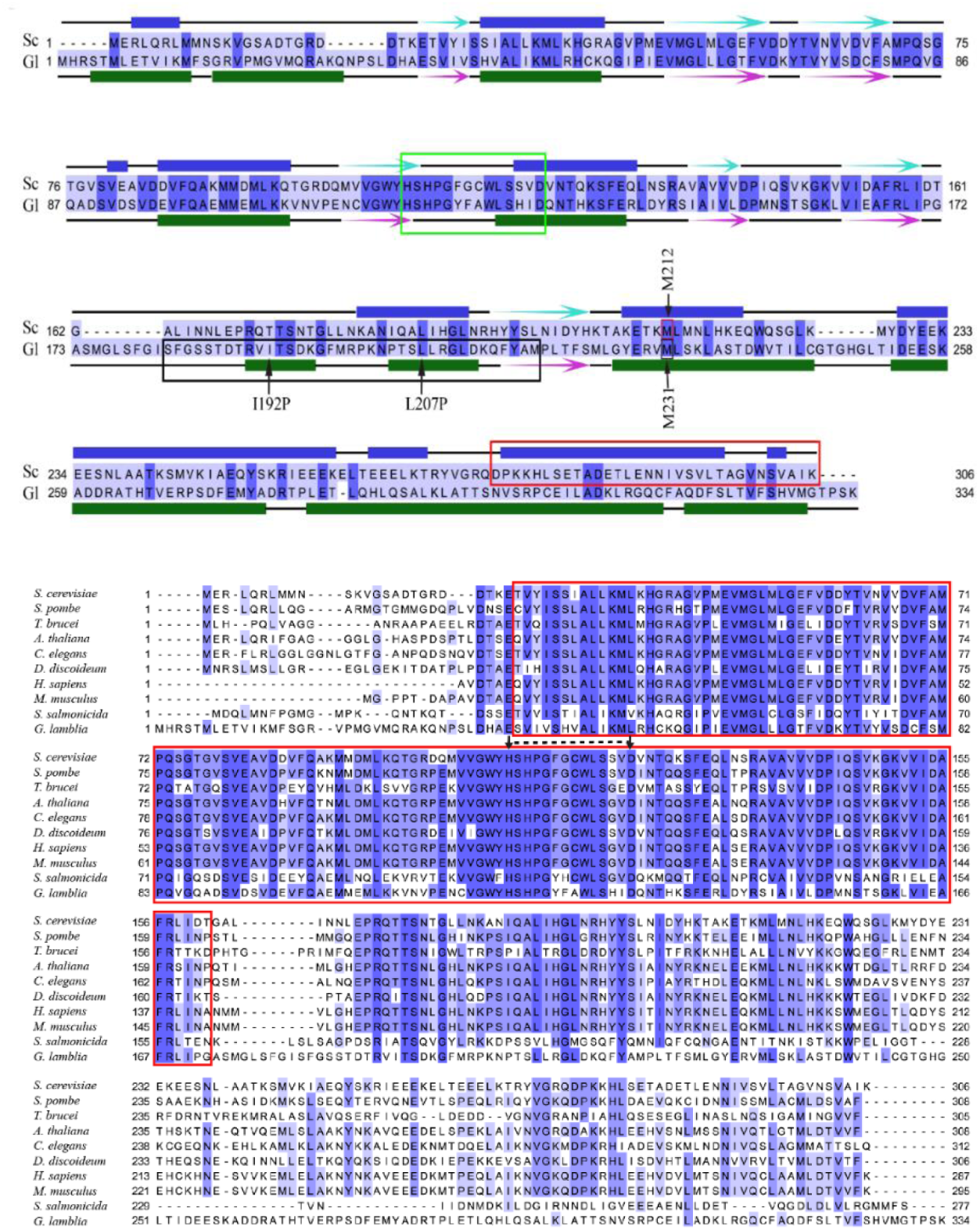
Multiple and pairwise sequence alignment of Rpn11. (**A**) Pairwise sequence alignment of Rpn11 from *Saccharomyces cerevisiae* (Sc) and *Giardia lamblia* (Gl) marking the position of conserved MPN+ motif (green box), the critical region of Rpn11 responsible for maintenance of mitochondrial structure and function (red box) and the functionally-equivalent region of GlRpn11 (black box). Residues Ile192 and Leu207 of GlRpn11 are marked with arrows. Met212 of Rpn11 and Met231 of GlRpn11 are also marked. The secondary structure elements of each protein, as predicted by Phyre^2^, are given either above (Sc) or below (Gl), where filled boxes represent α-helices, arrows represent β-sheets, and black lines represent unstructured regions. (**B**) Multiple sequence alignment of Rpn11 from different taxonomic groups (*Saccharomyces cerevisiae, Schizosaccharomyces pombe, Trypanosoma brucei, Arabidopsis thaliana, Caenorhabditis elegans, Dictyostelium discoideum, Homo sapiens, Mus musculus, Spironucleus salmonicida,* and *Giardia lamblia*) where the red box marks the position of the MPN domain and the black dashed line marks the position of the conserved MPN+ motif for all orthologues.

**Supplementary Figure 5.**
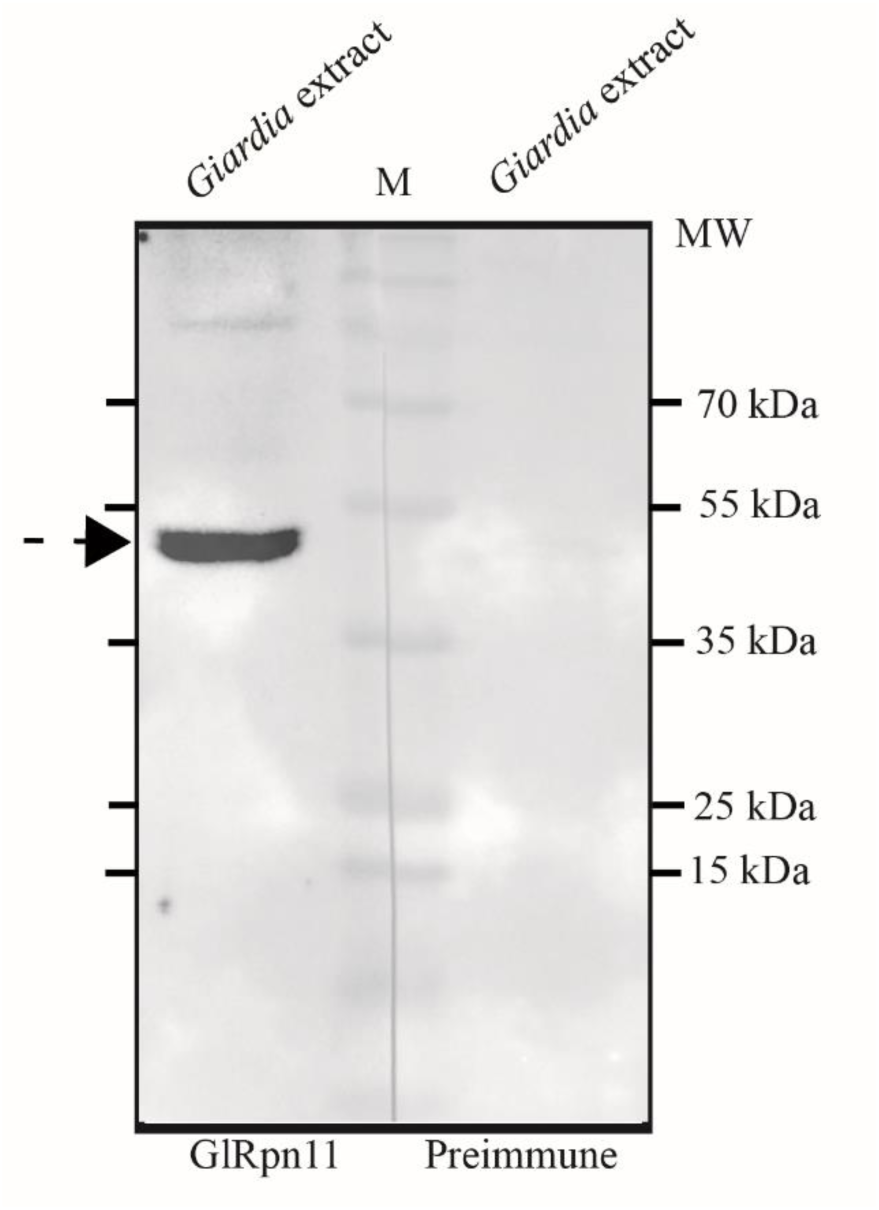
Western blot with an antibody against GlRpn11. Western blotting of *G. lamblia* trophozoite extract, with anti-GlRpn11; pre-immune sera served as control. The observed band of ∼ 36 kDa is consistent with the predicted size of GlRpn11.

**Supplementary Figure 6.**
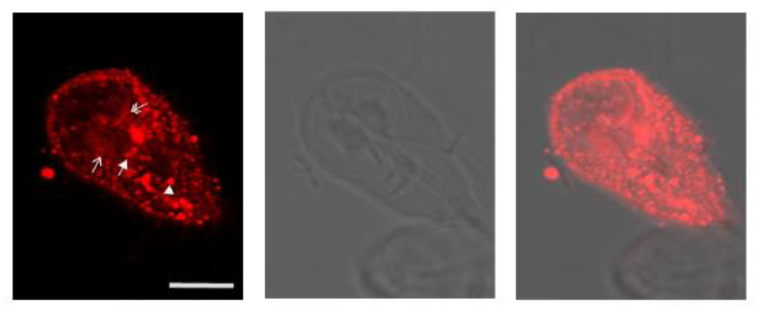
Association of GlRpn11 with ESV-like structures in encysting trophozoites. Immunofluorescence localization of GlRpn11 in 8 h post-induction of encystation. Alexa Flour 594 conjugated anti-rabbit IgG secondary antibody was used. The single-headed arrow indicates nuclear localization, the double-headed arrow indicates localization at the overlap zone of the ventral disc, the solid arrow marks the presence of GlRpn11 at the ventral flagella pore, and the solid arrowhead marks the presence the at structures whose size corresponds to those of encystation specific vesicles. The scale bar indicates 5 µm.

**Supplementary Figure 7.**
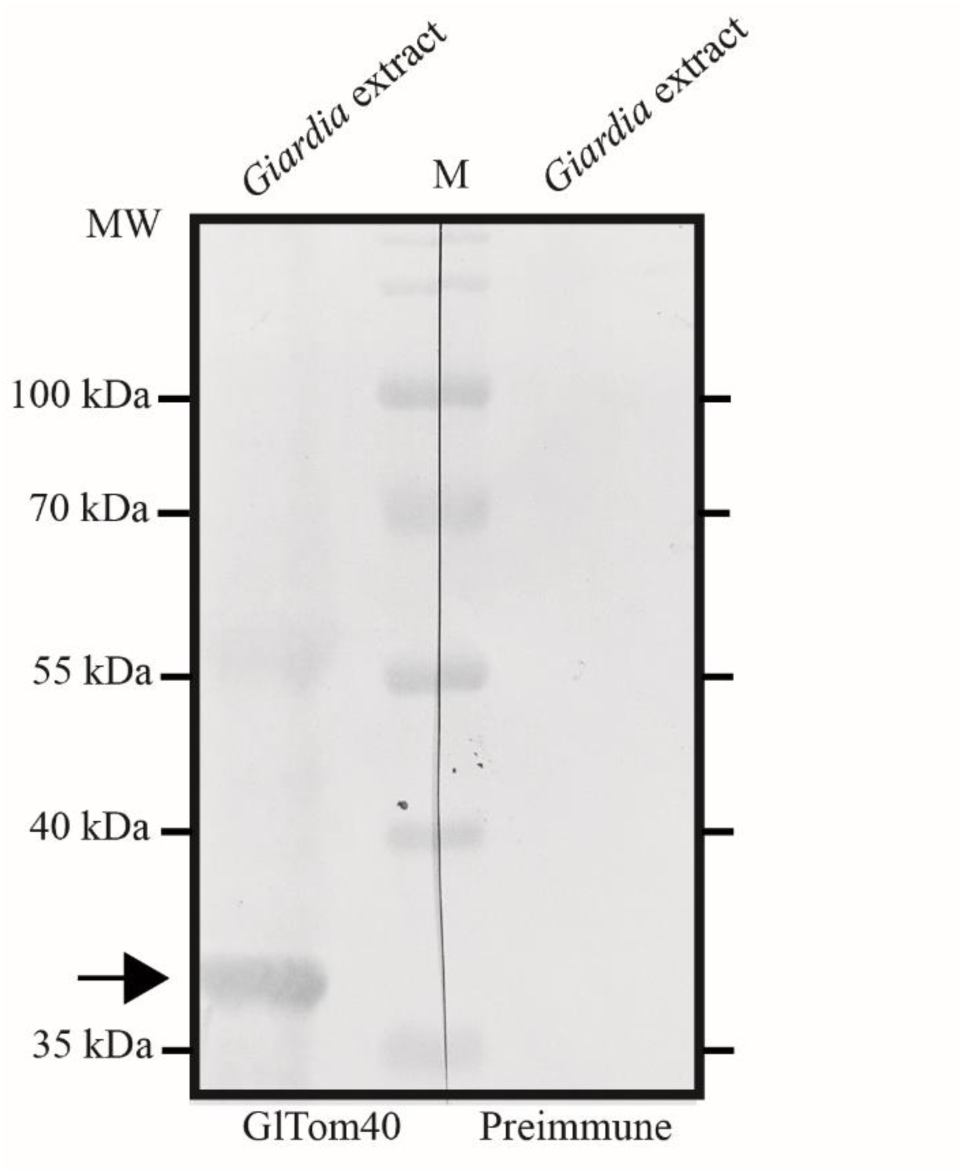
Western blot with an antibody against GlTom40. Western blotting of *G. lamblia* trophozoite extract, with anti-GlTom40; pre-immune sera served as control. The observed band of ∼ 39 kDa is consistent with the predicted size of GlTom40.

**Supplementary Figure 8.**
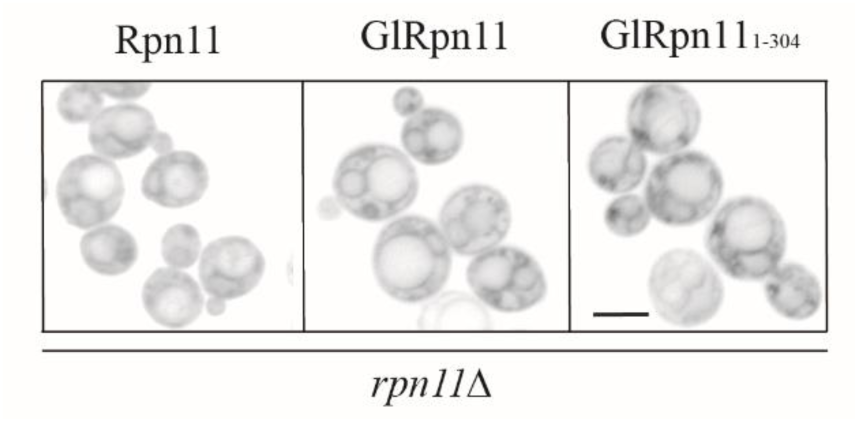
Mitochondrial network morphology of *rpn11Δ* cells expressing Rpn11, GlRpn11 or GlRpn11_1-304_. The morphology of the mitochondrial network of each strain, as indicated in the figure, was monitored using Mitotracker Deep Red FM. Rpn11 was expressed from a *URA3* vector, whereas GlRpn11 and GlRpn11_1-304_) were expressed from *LEU2* vector. The scale bar indicates 5 µm.

## Footnotes

1 3-AT, 3-amino-1,2,4-triazole; AD, Gal4 activation domain; BD, Gal4 DNA binding domain; CP, Core particle; LP, Lid particle; MSA, Multiple sequence alignment; OCR, Oxygen Consumption Rate; OZ, Overlap Zone; RP, Regulatory particle; VD Ventral Disc; VFP, Ventral Flagellar Pores; YCM, Yeast complete medium; Y2H, Yeast two-hybrid analysis

## Notes

### Competing Interest Statement

The authors have declared no competing interest.

